# Azelaic Acid as a Novel Antivirulence Agent Against *Pseudomonas aeruginosa*: In silico, In vitro and In vivo Efficacy and Biocompatibility

**DOI:** 10.64898/2026.03.18.712801

**Authors:** Maciel Espinosa Arriaga, Ana Paula Palacios Rodriguez, Rodolfo García Contreras, Geovany Esther Hu Oxte, Maricruz Sánchez Magaña, Gabriel Martínez González, Alejandra Ramírez Villalva, Jorge Almeida

## Abstract

*Pseudomonas aeruginosa* is an opportunistic pathogen whose virulence is largely regulated by quorum sensing (QS), making this system an attractive target for anti-virulence therapies. Azelaic acid (AzA), a naturally occurring dicarboxylic acid widely used in dermatology, has not previously been investigated as a QS inhibitor. This study evaluated the antibacterial and anti-virulence activities of AzA against reference strains and multidrug-resistant clinical isolates of *P. aeruginosa*, together with its preliminary safety profile and potential molecular interactions with key QS proteins. Minimum inhibitory concentrations (MICs) ranged from 250 to 1000 µg/mL. At sub-inhibitory concentrations, AzA significantly reduced the production of pyocyanin, elastase, protease and alginate in a dose-dependent manner without affecting bacterial viability. Molecular docking predicted favorable interactions of AzA with LasI, LasR, PqsD and PqsR, supporting a potential modulation of QS-associated pathways. Chemoinformatic analyses predicted favorable drug-likeness and safety characteristics, while PBMC viability remained above 90% and hemolysis below 4% within the anti-virulence concentration range. In the *Tenebrio molitor* infection model, AzA significantly increased larval survival and reduced in vivo pyocyanin production in both the reference strain Pa14 and a multidrug-resistant clinical isolate. Collectively, these findings provide the first evidence that AzA attenuates QS-regulated virulence in *P. aeruginosa* through complementary in vitro, in silico and in vivo approaches while exhibiting a favorable preliminary safety profile. These results support the repositioning of AzA as a promising anti-virulence candidate for the development of adjunctive therapeutic strategies against multidrug-resistant *P. aeruginosa* infections.

## 1. Introduction

Antimicrobial resistance (AMR) has emerged as one of the most pressing global public health threats of the 21st century. In the absence of preventive measures, projections indicate that by 2050, AMR could potentially supersede all other causes of mortality worldwide (1). Dolecek and colleagues discuss the GRAM report, published earlier this year, which estimated that bacterial AMR contributed to over 4 million deaths and was directly responsible for over 1.2 million deaths in 2019 (2). It was estimated that antibiotic resistance may have caused 1.27 million deaths in 2019, with 929,000 deaths attributable to the six leading pathogens*: Escherichia coli, Staphylococcus aureus, Klebsiella pneumoniae, Streptococcus pneumoniae, Acinetobacter baumannii,* and *Pseudomonas aeruginosa* (3).

It is important to pay special attention to WHO priority pathogens, including carbapenem-resistant *Enterobacteriaceae*, methicillin-resistant *Staphylococcus aureus* and extended-spectrum β-lactamase producers such as *E. coli and Klebsiella* spp., as well as multidrug-resistant *Pseudomonas aeruginosa* and carbapenem-resistant *Acinetobacter baumannii*, as these bacteria cause significant morbidity and mortality worldwide (4,5).

Among these, *Pseudomonas aeruginosa* stands out as a highly relevant threat due to its leading role in severe healthcare-associated infections, particularly nosocomial pneumonia, infections in immunocompromised hosts, and chronic colonization in patients with structural lung diseases like cystic fibrosis (6–8). The clinical success of *P. aeruginosa* relies heavily on its remarkable capacity to adapt to adverse host environments. This adaptability is driven by the synchronized secretion of a diverse array of virulence factors, which actively promote tissue damage, immune evasion, and the establishment of persistent infections (9).

*Pseudomonas aeruginosa*, uses chemical signals to communicate between cells in a process called quorum sensing (QS). QS allows groups of bacteria to sense population density and, in response to changing cell densities, to coordinate behaviors (10). It presents a particularly fine-tuned and convoluted network in which at least three interconnected systems Las, Rhl and Pqs perform in a hierarchical manner and a Pseudomonas quinolone signaling system 2-heptyl-3-hydroxy-4-quinolone (PQS)(11,12). LasI synthesizes the autoinducer N-(3-oxododecanoyl)homoserine lactone (OdDHL). OdDHL molecules bind to and activate the receptor protein LasR, ultimately triggering the transcription of virulence genes, such as the protease LasA, elastase LasB, alkaline protease, and exotoxin A (13). In the rhl system, the transcriptional activator RhlR recognizes N-butyryl-L-homoserine lactone (BHL), which is synthesized by the synthase RhlI (14). This system is tightly regulated by the las hierarchy at both transcriptional and post-translational levels (15). *Pseudomonas* quinolone signaling 2-heptyl-3-hydroxy-4-quinolone (PQS) has diverse functions, such as regulating the expression of virulence factors, iron acquisition, induction of oxidative stress and an antioxidant response, as well as modulating host immune responses(16).

Among the new therapeutic strategies to combat antibiotic resistance, antivirulence therapy has emerged as a promising alternative because, instead of killing pathogens, it attempts to deprive them of their virulence factors (17). QS inhibition is one of many strategies implemented to control microbial pathogenesis to reduce virulence and disarm pathogens rather than kill them. It is also attractive compared to other methods because it does not impose strong selective pressure and, therefore, bacterial resistance is less likely to develop (18,19). Therefore, QSIs provide an exceptional advantage over conventional chemical antibiotics and disinfectants, which exert selective pressure through their bactericidal action (20). In this context, azelaic acid (AzA), a saturated dicarboxylic acid that exhibits antibacterial activity and recent research indicates that AzA can also inhibit biofilm formation by *C. acnes*, reducing bacterial virulence, making it a molecule to explore for its QSI activity.

Therefore, the aim of this study was to evaluate the antimicrobial activity of azelaic acid (AzA) by determining the minimum inhibitory concentration (MIC) against seven reference strains—*P. aeruginosa* (Pa01, Pa14, and ATCC 27853), *E. coli* (ATCC 25922), *Proteus mirabilis* (ATCC 43091), methicillin-resistant *Staphylococcus aureus* (MRSA, ATCC 43300), *S. aureus* (ATCC 6538), and *K. pneumoniae* (ATCC 33495)—and four clinical isolates, including an ESBL-producing *E. coli* strain, a clinical *S. aureus* strain, and two resistant *P. aeruginosa* strains (one PDR and one XDR). Furthermore, this research focused on the anti-virulence potential of AzA by evaluating the inhibition of quorum sensing-regulated factors—specifically the production of pyocyanin, elastase, alginate, and exoproteases—in the five *P. aeruginosa* strains. To gain insight into the mechanism of action, molecular docking simulations were performed to predict the interaction of AzA with key quorum sensing receptor proteins. As this study involved drug repurposing, safety was assessed through cytotoxicity tests on PBMCs and hemolysis assays; additionally, survival in an infected, treated model was evaluated using Kaplan-Meier curves, and *in vivo* pyocyanin quantification was performed to confirm that virulence attenuation occurs not only *in vitro* but also *in vivo*. To the best of our knowledge, this is the first study to describe the inhibitory activity of azelaic acid on quorum sensing in *Pseudomonas aeruginosa*, offering a novel perspective on its potential as an anti-virulence agent.

## 2. Materials and Methods

### 2.1. Preparation of strains and minimum inhibitory concentration

The study included a diverse panel of reference and clinical strains. The reference strains consisted of *P. aeruginosa* (Pa01, Pa14, and ATCC 27853*), S. aureus* (ATCC 43300 and ATCC 6538), *E. coli* (ATCC 25922*), K. pneumoniae* (ATCC 33495*), P. mirabilis* (ATCC 43091). Additionally, four clinical strains were evaluated: PaHer, a multidrug-resistant (MDR) strain obtained from a wound with cephalosporin resistance; PaBro, a pandrug-resistant (PDR) strain isolated from bronchial secretion; *E. coli* Ur, showing an extensively drug-resistant (XDR) and extended-spectrum β-lactamase (ESBL) pattern; and a susceptible *S. aureus* Ur strain, both obtained from urine cultures. The identification of clinical strains as well as their antibiotic resistance patterns was carried out using the automated Microscan WalkAway method. These strains were incubated on BHI and LB agar plates for *Pseudomonas* for at least 20 hours at 37°C to activate the culture.

### 2.2. Minimum Inhibitory Concentration (MIC) Determination

The minimum inhibitory concentration (MIC) was determined using the microdilution method in a 96-well plate (Nunc), according to the guidelines in Table 8A of the Clinical and Laboratory Standards Institute (CLSI) (21,22). Azelaic acid (AzA) was dissolved in DMSO and tested at concentrations ranging from 62.5 to 1000 µg/mL. The bacterial inoculum was adjusted to approximately 10⁸ CFU/mL of microorganisms. Subsequently, 10 µL of the inoculum were added to wells containing Mueller Hinton Broth (MHB), achieving a final concentration of 10⁵ CFU/mL (23). Control groups included: MHB without microorganisms, MHB with the strains (growth control), and MHB with DMSO and strains to confirm that it did not interfere with bacterial growth. The microplates were incubated at 35 ± 2°C for 20 h. All assays were performed in quadruplicate.

### 2.3. Strain preparation and treatment with azelaic acid

To activate the *P. aeruginosa* strains (Pa01, Pa14, ATCC 27853, PaBro, and PaHer), these were subcultured on LB agar plates (1.5% agar) and incubated for 20 h at 37 °C. Subsequently, overnight (ON) cultures were prepared by inoculating each strain into 50 mL flasks containing 25 mL of LB medium, followed by incubation for 18 h at 37 °C. Bacterial growth was then monitored by measuring the optical density at 600 nm (OD_600_) using a 1:10 dilution in LB broth.

Fresh cultures were then inoculated from the ON stocks to a standardized OD_600_. For each of the five strains, three series of flasks (each containing 25 mL of LB medium) were prepared to evaluate different concentrations of azelaic acid (AzA): 250, 500, and 750 µg/mL (sub-MIC concentrations of 1/4, 1/2, and 3/4 respectively). The AzA was dissolved in DMSO and sterilized using 0.22 µm syringe filters prior to addition.

The cultures were incubated at 37 °C with constant shaking for 24 h. Growth was monitored at OD_600_ for both treated samples and controls. Controls containing only DMSO were included to ensure the solvent did not interfere with bacterial growth, and a medium-only control was used to establish the baseline growth.

### 2.4. Determination of Pyocyanin

Aliquots (1mL) from the overnight cultures were centrifuged at 13,000 rpm for 1.5 min. A volume of 800µL of the supernatant was collected and mixed with 420µL of chloroform. The mixture was vortexed for 1 min and centrifuged at 13,000 rpm for 5min.

Subsequently, 300 µL of the organic (bottom) phase was transferred to a new tube containing 800 µL of 0.1N HCl. The solution was vortexed for 1 min and centrifuged at 13,000 rpm for 5 min to facilitate phase separation. Pyocyanin concentration was determined by measuring the absorbance at 520 nm using a quartz cuvette (24).

### 2.5. Protease Determination

To evaluate extracellular protease production, 1mL of the overnight cultures were centrifuged at 13,000 rpm for 2 min. A reaction mixture was prepared by combining 875µL of protease substrate buffer with 125 µL of the culture supernatant. After vortexing for 1 min, the samples were incubated at 37°C with shaking at 200 rpm for 1h. The reaction was stopped by adding 1% HNO_3_ and vortexing for 1 min. The mixture was then centrifuged at 13,000 rpm for 2 min. The resulting supernatant was mixed with 0.5% NaOH and gently stirred. Finally, the samples were transferred to quartz cuvettes, and the absorbance was measured at 595 nm (25).

### 2.6. Determination of alginate

To determinate the production of alginate, 1mL of the overnight cultures were centrifuged and subsequently heated at 80° C for 30 min. The resulting supernatant was centrifuged for 30 min at 13,000 rpm, the pellet was removed, and the alginate was precipitated with ice-cold 99% ethanol. The precipitated alginate was collected and dissolved in 1 mL of 0.9% sterile saline. A reaction mixture was prepared by combining 118 µL of sample with 1 mL of H_3_BO_3_/H_2_SO_4_ and 34 µL of carbazole. This mixture was heated at 55°C for 30 min and the absorbance was measured at 530 nm (OD _530_) (26).

### 2.7. Determination of Elastase

1mL aliquot was collected from each culture and centrifuged at 13,000 rpm for 1.5 minutes, after which the cell-free supernatant was recovered. Subsequently, 100 µL of the supernatant from each culture were mixed with 5 mg of elastin-Congo red (Sigma-Aldrich) and 900 µL of Tris-HCl buffer (pH 7.5). The mixture was incubated at 37 °C for 2 h and then centrifuged at 13,500 rpm for 5 minutes. Finally, the absorbance was measured at 495 nm (27).

### 2.8. Statistical analysis

An analysis of variance (ANOVA) and Tukey HSD test were performed to evaluate the effect of azelaic acid treatment at three different concentrations (250 µg/ml, 500 µg/ml, and 750 µg/ml) on the bacterial strains Pa01, Pa14, ATCC27853 PaHer, and PaBro. The ANOVA analysis, Tukey HSD test, and standard deviation (SD) calculations were performed using IBM SPSS Statistics for Windows, version 21.0 (IBM Corp., Armonk, NY).

### 2.9. Bioinformatic analysis

#### 2.9.1. Molecular and chemoinformatic properties

The molecular and cheminformatics properties were evaluated using the pkCSM, Molsoft, ChemSketch, SwissADME, and Osiris Property Explorer simulators. The values obtained for Azelaic acid are presented in Table 3. According to Lipinski’s rules, a compound with good oral bioavailability should meet the following criteria: LogP < 5, topological polar surface area (TPSA) < 140 Å², molecular weight < 500 Da, fewer than 5 hydrogen bond donors, and fewer than 10 hydrogen bond acceptors. Compliance with these parameters suggests favorable therapeutic potential.

#### 2.9.2. Ligand preparation

Azelaic acid was selected as the test ligand and retrieved from the PubChem database in SDF format. Ligand preparation was performed using OpenBabel GUI, where the molecule was deprotonated to its dianionic form to reflect the predominant ionization state at a physiological pH of 7.4 (pKa₁ ≈ 4.5, pKa₂ ≈ 5.4). The geometry was subsequently optimized using the MMFF94 force field in Avogadro2 to obtain a stable conformation. The optimized structure was then converted to MOL2 format, and Gasteiger charges were assigned using AutoDock Tools v. 1.5.6 in preparation for docking calculations.

#### 2.9.3. Protein receptor preparation

X-ray crystallographic structures of proteins involved in the quorum sensing system of *Pseudomonas aeruginosa* were obtained from the Protein Data Bank (PDB), including LasI (PDB ID: 1RO5), LasR (PDB ID: 3IX3 chain A and 2UV0 chain E), PqsD (PDB ID: 3H76), and PqsR (PDB ID: 4JVI). Protein preparation was conducted in ChimeraX, where water molecules, heteroatoms, and co-crystallized ligands were removed. Native ligands, when present, were extracted for protocol validation. Polar hydrogens were added, and Kollman charges were assigned using AutoDock Tools v.1.5.6, after which the structures were converted to PDBQT format.

#### 2.9.4. Molecular docking

Docking calculations were performed using AutoDock Vina 1.1.2 with default parameters (exhaustiveness = 8, maximum number of binding modes = 9, energy range = 3 kcal/mol). For proteins with co-crystallized ligands, protocol validation was performed by redocking the native ligand into its original binding site, and docking accuracy was evaluated via root mean square deviation (RMSD), with protocols considered valid when RMSD ≤ 2.0 Å. For LasI (PDB ID: 1RO5), which lacks a co-crystallized ligand, a focused docking approach was applied based on the reported active-site region, using TZD-C8 (Z-5-octylidene-thiazolidine-2,4-dione) as a positive control to confirm the docking setup (28–30). For PqsD (PDB ID: 3H76), only literature references were used to define the docking grid (31).

Following validation, azelaic acid was docked into each receptor using grid boxes centered on either the crystallographic ligand or reported active-site regions. The grid box coordinates and dimensions were as follows: 1RO5 (x: 47.179, y: −9.516, z: −9.139; size x: 20.25, y: 18.75, z: 21.75), 3IX3 (x: 10.54, y: 4.32, z: 20.98; size x: 18, y: 18, z: 18), 2UV0 (x: 24.8464, y: 14.42, z: 78.965; size x: 18, y: 18, z: 18), 3H76 (x: −9.917, y: −7.196, z: −3.603; size x: 21, y: 22.5, z: 22.5), and 4JVI (x: −34.222, y: 56.883, z: 9.326; size x: 18, y: 18, z: 18).

The best-ranked ligand poses were selected based on predicted binding affinity (kcal/mol) and visual inspection of key interactions within the active site. Protein–ligand interactions, including hydrogen bonds, salt bridges, hydrophobic contacts, and π interactions, were further characterized using Discovery Studio Visualizer and the Protein-Ligand Interaction Profiler (PLIP) (32). The use of PLIP provided a rigorous cross-validation of ionic interactions and bond distances, ensuring the chemical accuracy of the predicted binding modes. This workflow ensured consistent and reproducible docking results across all evaluated receptors.

### 2.10 In vivo evaluation using Tenebrio molitor

The Pa14 and PaBro strains were standardized by culturing overnight in LB broth to an OD600nm corresponding to 5 × 10⁵ CFU in PBS. Of this preparation, 10 µL were injected into *Tenebrio molitor* using a microsyringe needle (33). This served as an infection control for comparison against the treatment. Inoculation was performed through the pleural intersegmental membrane between abdominal segments 2 and 4, ensuring the safest route for the pathogen to enter the hemocoel directly without causing physical trauma(34,35). The following were used: an untreated, non-inoculated control; a PBS control; treatments with AzA at 100, 250, 500, and 750 µg/mL; and—to further evaluate AzA toxicity—a control at 1000 µg/mL without the strain. After injection, the larvae were incubated in a Petri dish at 30°C for 3 days. Larval survival rates were determined by counting the number of dead larvae every 12 hours (36). The Pa14 strain was selected as the most virulent of the reference strains, and the PaBro strain was chosen for its resistance pattern and to determine whether the effect observed in vitro would translate to in vivo conditions. To evaluate and compare the impact of infection by both strains and the efficacy of treatment at different concentrations in the biological model, survival curves were constructed using the non-parametric Kaplan-Meier estimator. The event of interest was defined as the death of the *Tenebrio molitor* larva, with survival time recorded in days from the moment of inoculation (t0). Differences in survival rates among the untreated infected groups and the treated infected group—as well as PBS controls, untreated/uninfected controls, and a safety control using only azelaic acid at 1000 µg/mL—were statistically evaluated using the log-rank (Mantel-Cox) test, with a p-value < 0.05 considered statistically significant (37).

### 2.11 PBMC Cytotoxicity and Hemolysis

#### 2.11.1 Hemolysis

To quantitatively determine cytotoxic effects on human erythrocytes, a hemolytic activity test was performed using a sample of approximately 4 mL of human peripheral blood. Two milliliters of blood were transferred to a 15 mL conical tube and centrifuged at 1500 rpm for 5 minutes at 30°C. The resulting plasma was removed, and PBS (Phosphate-Buffered Saline) was added; the mixture was homogenized two to three times using a micropipette and centrifuged again under the previously described conditions (this process was performed in triplicate). Upon completion, the supernatant was removed, and 80 µL of the erythrocyte pellet was collected and mixed with 3920 µL of PBS to obtain a 2% erythrocyte solution.

Subsequently, 100 µL of the 2% erythrocyte solution was added to each well of a 96-well plate, followed by the addition of appropriate amounts of azelaic acid to achieve final concentrations of 750, 500, 300, 200, 100, 10, and 1 µg/mL. For the positive control, 100 µL of 1% SDS was added to the 2% erythrocyte solution; for the negative control, 100 µL of the 2% erythrocyte solution was mixed with 100 µL of PBS. The plate was incubated at room temperature for three hours, with the contents being resuspended every 60 minutes. After the incubation period, the plate was centrifuged at 1500 rpm for 5 minutes at 37°C. Finally, 100 µL was taken from each well—ensuring no erythrocytes were drawn up—and transferred to a new 96-well plate for reading at 340 nm using an iMark Microplate Reader. To obtain the percentage of hemolysis, the calculation was performed using the following formula (38):

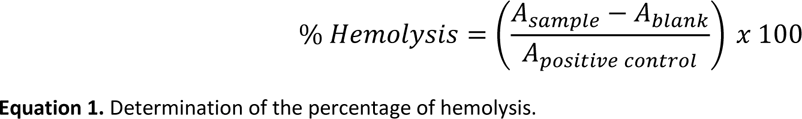

#### 2.11.2 MTT viability assay on PBMCs

For the MTT cytotoxicity assay, a 4 mL peripheral blood sample was diluted 1:1 with PBS, and 3 mL of this mixture was transferred to a 15 mL tube (Falcon™) containing 3 mL of Ficoll. The sample was centrifuged at 400 × g for 40 minutes. The resulting buffy coat was collected in a 15 mL tube, and 6 mL of PBS was added for a wash step, followed by centrifugation at 500 × g for 10 minutes. The supernatant was decanted, and a second wash was performed under the same conditions. After this step, the supernatant was removed, and the cells were resuspended in 1 mL of DMEM medium supplemented with FBS. A 1:10 dilution of this suspension with trypan blue was prepared in an Eppendorf tube; a 10 µL aliquot was then loaded into a Neubauer chamber for microscopic observation and cell counting, and the concentration was adjusted to 50,000 cells per well (requiring a total of 3,000,000 cells). Subsequently, 100 µL of the cell suspension was added to each well, and the plate was incubated for 24 hours at 37°C in a 5% CO_2_ atmosphere. Following incubation, the supernatant was removed, and azelaic acid dissolved in DMEM medium was added in appropriate amounts to achieve final concentrations of 1000, 750, 500, 300, 200, 100, 10, and 1 µg/mL. DMSO (100 µL) was used as the positive control, and DMEM medium (100 µL) was used as the negative control. The plate was incubated for 24 hours at 37°C with 5% CO_2_. After the incubation period, the plate was centrifuged at 1500 rpm for 5 minutes. Following exposure of the cells to the different concentrations, the supernatant was removed to perform the viability assay using the MTT colorimetric reaction (at a concentration of 5 mg/mL); the tetrazolium salt was added to the pre-treated cells, and the plate was incubated for 4 hours. After this time, the supernatant was removed, and DMSO reagent was added to each well. The plate was allowed to stand for 10 minutes, and absorbance was measured using a plate reader at 570 nm. High absorbance values correlated with higher total metabolic activity and a greater number of viable cells. The obtained absorbance values were converted into a percentage of cytotoxicity using the following formula(39):

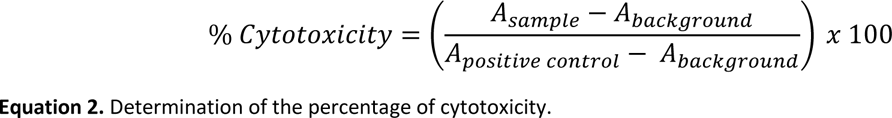

### 2.12 *In vivo* quantification of pyocyanin

To obtain pyocyanin *in vivo*, a liquid-liquid extraction method—originally classical but modified for complex solid matrices—was adapted. Since the pigment is distributed throughout both the hemolymph and the colonized internal tissues of *Tenebrio molitor* (40), mechanical lysis via maceration was essential to ensure the complete release of the bacterial metabolite. Subsequently, the crude extract was centrifuged at 13,000 rpm for 20 minutes, causing insoluble chitin residues and cellular debris to precipitate into the pellet. The particle-free supernatant was recovered as the initial aqueous phase, thereby ensuring optimal interaction—free of emulsions—with the chloroform during the subsequent pH-gradient separation steps (41). The remainder of the pyocyanin extraction and quantification was performed as described in section 2.4.

## 3. Results

### 3.1. Antimicrobial Susceptibility (MIC)

The antibacterial activity of azelaic acid (AzA) was determined using the broth microdilution method. For all strains tested, including reference pathogens and clinical isolates of *P. aeruginosa* (PaBro “PDR” and PaHer “MDR”), exhibited a Minimum Inhibitory Concentration (MIC) of 1000 µg/mL. In contrast, for clinical isolates of *E. coli* Ur and *S. aureus* Ur, the MIC was at 250 µg/mL (Table 1). Based on these results, subinhibitory concentrations of 250, 500, and 750 µg/mL (representing 1/4, 1/2, and 3/4 of the MIC for *P. aeruginosa*, respectively) were selected for subsequent quorum sensing inhibition assays to ensure that the observed effects were due to virulence attenuation and not bactericidal activity.

**Table 1.**
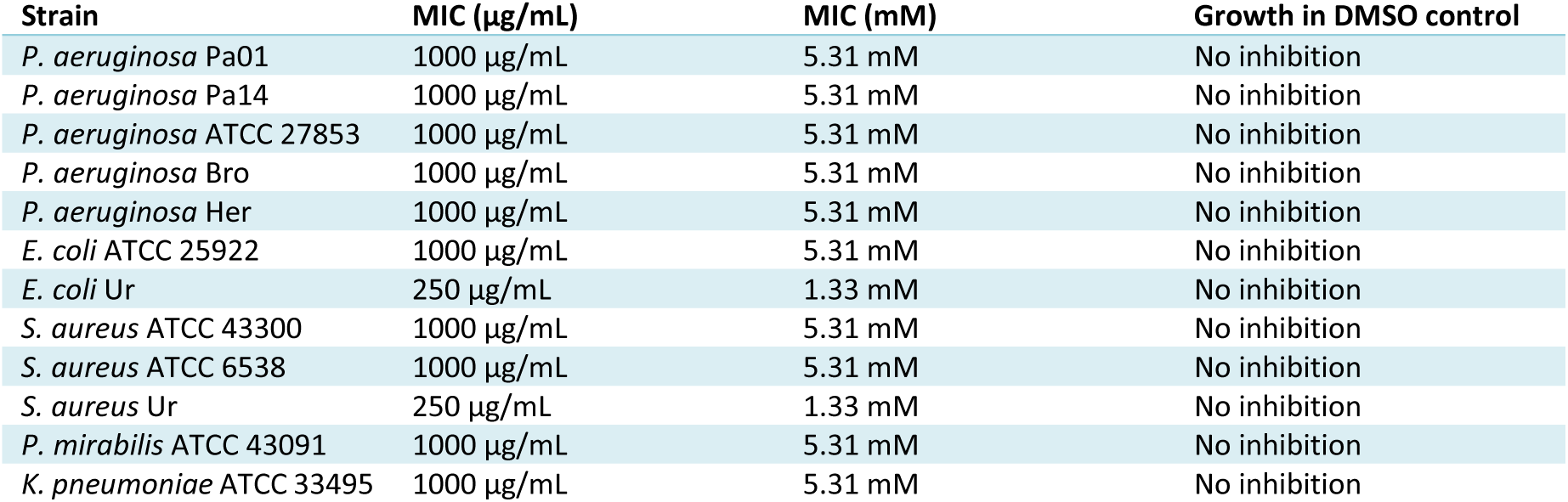
Minimum Inhibitory Concentration (MIC) of azelaic acid against various strains of bacteria.

| Strain | MIC (µg/mL) | MIC (mM) | Growth in DMSO control |
| --- | --- | --- | --- |
| *P. aeruginosa* Pa01 | 1000 µg/mL | 5.31 mM | No inhibition |
| *P. aeruginosa* Pa14 | 1000 µg/mL | 5.31 mM | No inhibition |
| *P. aeruginosa* ATCC 27853 | 1000 µg/mL | 5.31 mM | No inhibition |
| *P. aeruginosa* Bro | 1000 µg/mL | 5.31 mM | No inhibition |
| *P. aeruginosa* Her | 1000 µg/mL | 5.31 mM | No inhibition |
| *E. coli* ATCC 25922 | 1000 µg/mL | 5.31 mM | No inhibition |
| *E. coli* Ur | 250 µg/mL | 1.33 mM | No inhibition |
| *S. aureus* ATCC 43300 | 1000 µg/mL | 5.31 mM | No inhibition |
| *S. aureus* ATCC 6538 | 1000 µg/mL | 5.31 mM | No inhibition |
| *S. aureus* Ur | 250 µg/mL | 1.33 mM | No inhibition |
| *P. mirabilis* ATCC 43091 | 1000 µg/mL | 5.31 mM | No inhibition |
| *K. pneumoniae* ATCC 33495 | 1000 µg/mL | 5.31 mM | No inhibition |

### 3.2. Inhibition of Virulence Factors by Azelaic Acid (AzA)

AzA demonstrated a consistent, dose-dependent inhibitory effect on the production of all evaluated virulence factors. Control groups (untreated and DMSO) showed 0% inhibition, confirming that the effects were strictly attributable to AzA (table 2).

**Table 2.**
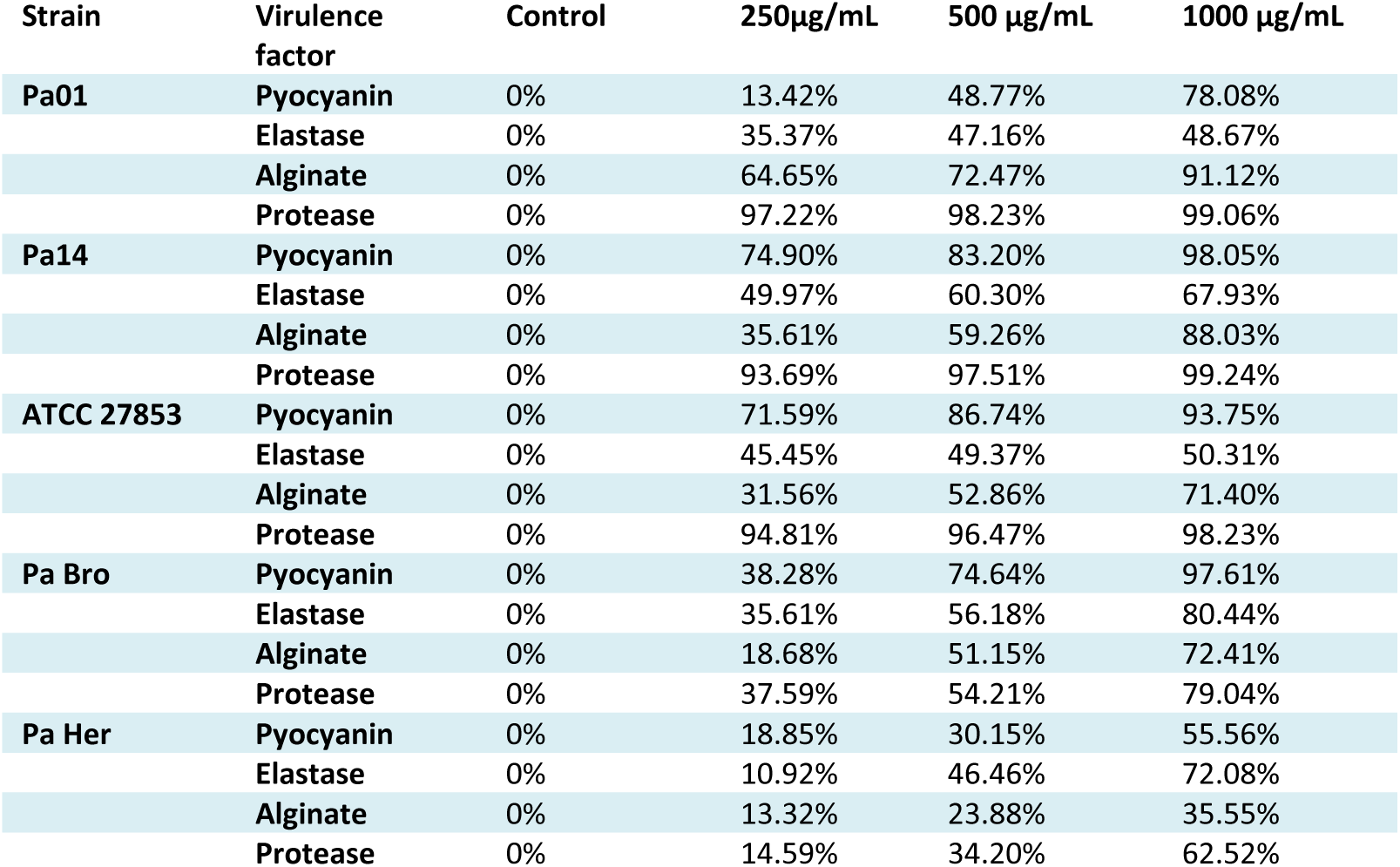
Percentage of inhibition of virulence factors of *P. aeruginosa* under treatment with azelaic acid.

| Strain | Virulence factor | Control | 250µg/mL | 500 µg/mL | 1000 µg/mL |
| --- | --- | --- | --- | --- | --- |
| **Pa01** | **Pyocyanin** | 0% | 13.42% | 48.77% | 78.08% |
|  | **Elastase** | 0% | 35.37% | 47.16% | 48.67% |
|  | **Alginate** | 0% | 64.65% | 72.47% | 91.12% |
|  | **Protease** | 0% | 97.22% | 98.23% | 99.06% |
| **Pa14** | **Pyocyanin** | 0% | 74.90% | 83.20% | 98.05% |
|  | **Elastase** | 0% | 49.97% | 60.30% | 67.93% |
|  | **Alginate** | 0% | 35.61% | 59.26% | 88.03% |
|  | **Protease** | 0% | 93.69% | 97.51% | 99.24% |
| **ATCC 27853** | **Pyocyanin** | 0% | 71.59% | 86.74% | 93.75% |
|  | **Elastase** | 0% | 45.45% | 49.37% | 50.31% |
|  | **Alginate** | 0% | 31.56% | 52.86% | 71.40% |
|  | **Protease** | 0% | 94.81% | 96.47% | 98.23% |
| **Pa Bro** | **Pyocyanin** | 0% | 38.28% | 74.64% | 97.61% |
|  | **Elastase** | 0% | 35.61% | 56.18% | 80.44% |
|  | **Alginate** | 0% | 18.68% | 51.15% | 72.41% |
|  | **Protease** | 0% | 37.59% | 54.21% | 79.04% |
| **Pa Her** | **Pyocyanin** | 0% | 18.85% | 30.15% | 55.56% |
|  | **Elastase** | 0% | 10.92% | 46.46% | 72.08% |
|  | **Alginate** | 0% | 13.32% | 23.88% | 35.55% |
|  | **Protease** | 0% | 14.59% | 34.20% | 62.52% |

### 3.3. Pyocyanin Production

As shown in Figure 1A, AzA significantly reduced pyocyanin secretion. The most notable effects were observed in the Pa14 and PaBro (PDR) strains, reaching inhibition levels of 98.05% and 97.61%, respectively, at 750 µg/mL. In contrast, the PaHer (MDR) clinical isolate showed a more moderate response, with a maximum inhibition of 55.56%. Analysis of variance (ANOVA) confirmed a highly significant effect of azelaic acid on all strains (p < 0.001). Tukey’s HSD test validated the dose dependence, showing significant differences (p < 0.001) between each concentration.

**Figure 1A.**
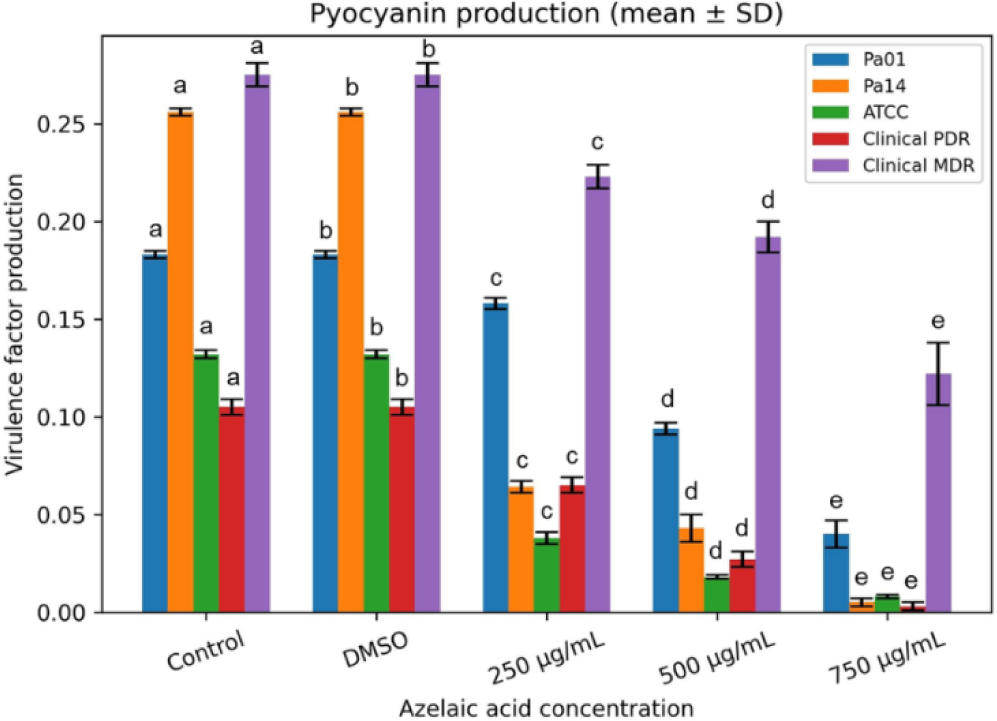
Inhibition of pyocyanin production by azelaic acid A dose-dependent reduction in pigment synthesis was observed in all strains studied. ANOVA analysis confirmed highly significant differences (p < 0.001).

### 3.4. Alginate Synthesis

The inhibitory effect of azelaic acid (AzA) on alginate production is illustrated in Figure 1B. Among the evaluated strains, the reference strain PaO1 was the most susceptible, exhibiting a 91.12% reduction at the highest concentration (750 µg/mL). The clinical isolates, PaBro (PDR) and PaHer (MDR), showed 72.41% and 35.55% inhibition, respectively, at the same dosage. These results suggest varying degrees of baseline mucoidity and metabolic resistance among the clinical strains.

**Figure 1B.**
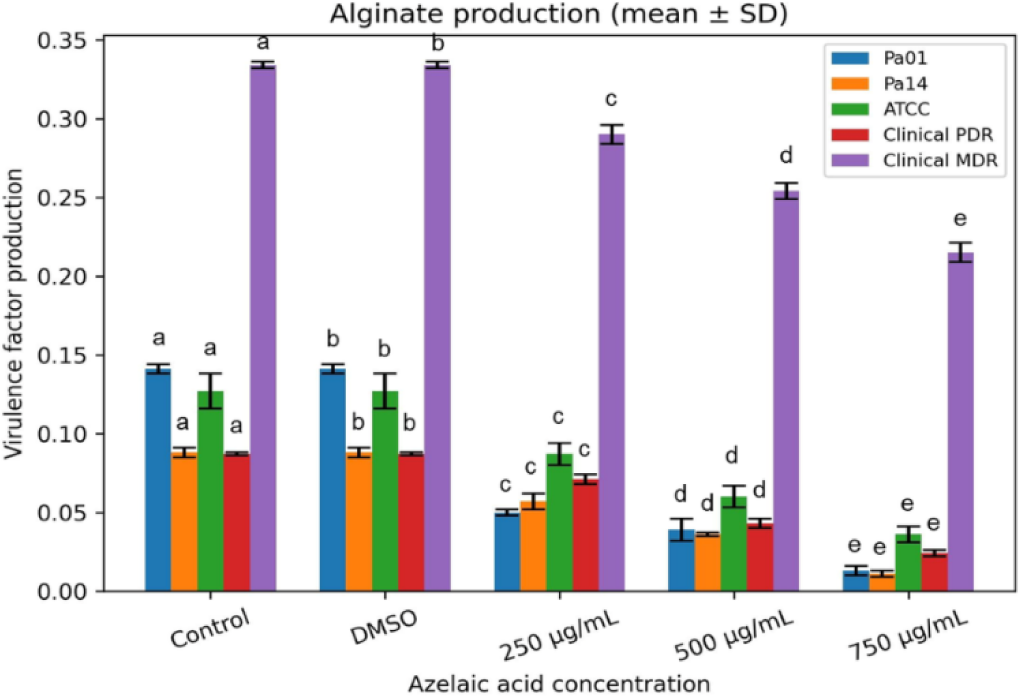
Effect of azelaic acid on alginate. The treatment significantly reduced the presence of this exopolysaccharide. The F-values demonstrate robust variance between groups, particularly in PaOl {F = 970.497} and the clinical strain Pa Bro {F = 538.547) (p < 0.001).

Analysis of variance (ANOVA) demonstrated a highly significant inhibitory effect of azelaic acid on alginate production across all strains (p < 0.001). Furthermore, Tukey’s HSD test confirmed the dose-dependent nature of this effect, validating significant differences between all concentrations tested.

### 3.5. Exoprotease Activity

As illustrated in Figure 1C, exoprotease production was the most sensitive virulence factor to AzA treatment. All reference strains achieved over 93% inhibition at the lowest concentration tested (250 µg/mL). The highest inhibitory activity in the entire study was recorded for the Pa14 strain at 750 µg/mL (99.24%).

**Figure 1C.**
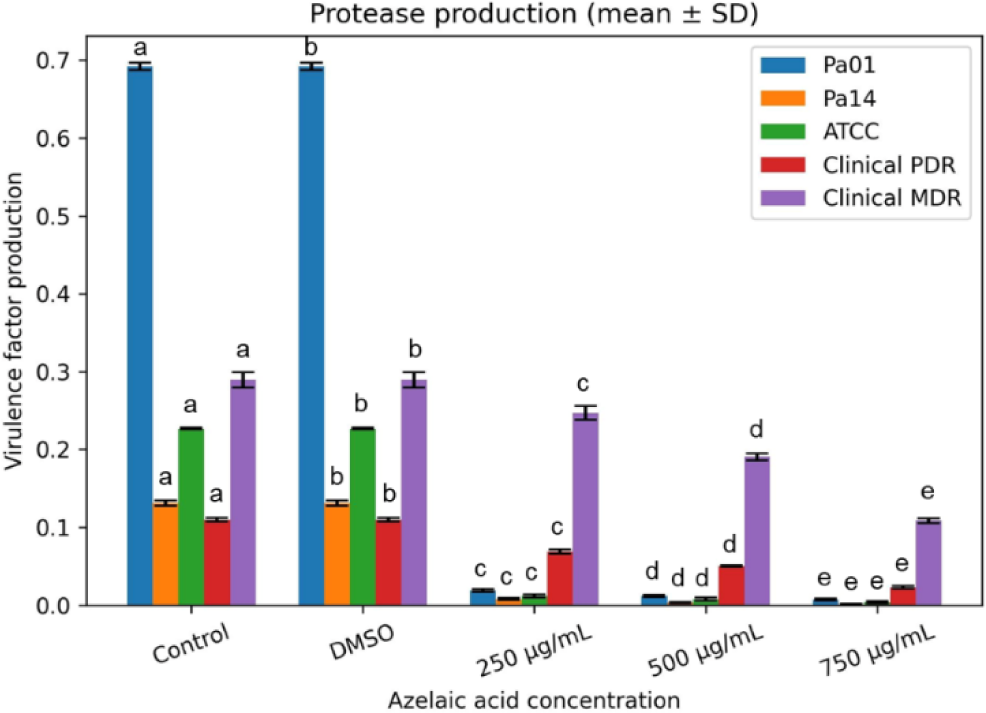
Suppression of proteolytic activity. Exposure to azelaic acid resulted in a drastic decrease in protease secretion, reaching inhibitions greater than 99% (p < 0.001).

Treatment with azelaic acid demonstrated robust inhibitory capacity on proteolytic activity in all strains evaluated (p < 0.001), supported by exceptionally high F values in the ANOVA analysis. The Tukey HSD post hoc test confirmed that this reduction is not only statistically significant compared to the control, but is also strictly concentration-dependent, achieving almost total inhibition (>99%) in the reference strains.

### 3.6. Elastase Inhibition

The reduction of elastolytic activity is presented in Figure 1D. Although elastase proved to be more resilient than general proteases, AzA achieved significant attenuation, particularly in the PDR clinical isolate (80.44%) and the Pa14 strain (67.93%) at 750 µg/mL.

**Figure 1D.**
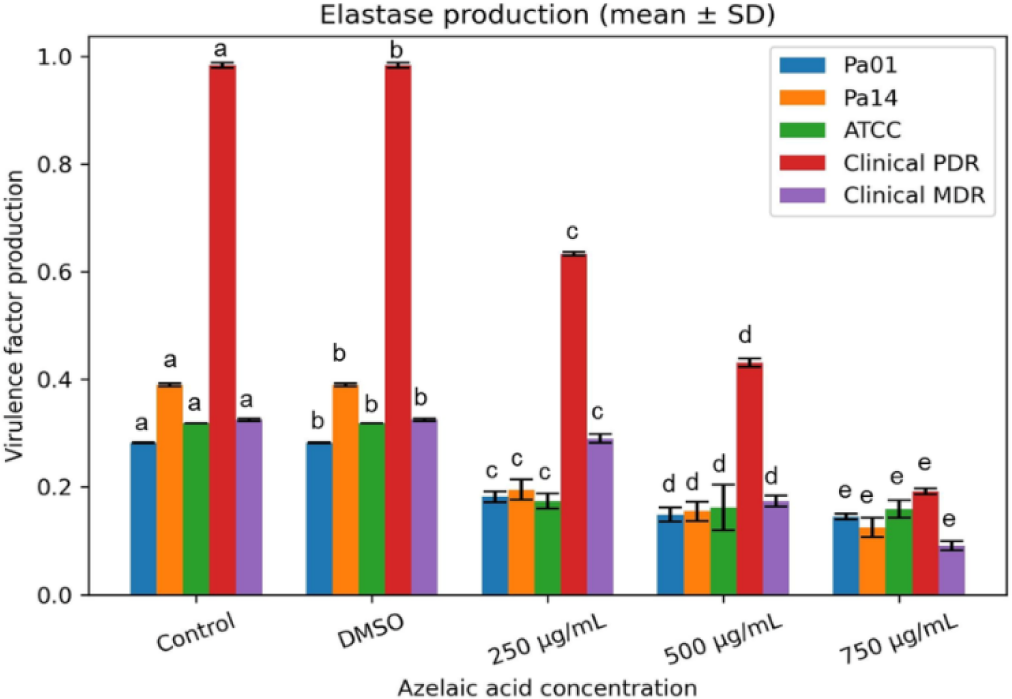
Elastolytic activity of P. aeruginosa. The graph illustrates the compound’s effectiveness in mitigating elastase production. The Tukey HSD post hoc test shows a progressive and statistically significant reduction (p < 0.001).

ANOVA results indicated that azelaic acid significantly mitigates elastase secretion across all studied strains (p < 0.001). Furthermore, Tukey’s HSD multiple comparisons analysis corroborated that this enzyme inhibition is strictly dose-dependent, achieving reductions of up to 80% in elastolytic activity.

Overall, azelaic acid showed an inhibitory effect on *Pseudomonas aeruginosa* virulence factors regulated by quorum sensing, including the production of elastase, protease, pyocyanin, and alginate. The highest concentration tested (750 μg/mL) produced substantial inhibition percentages in all phenotypes analyzed, with statistical significance confirmed by one-way ANOVA followed by Tukey’s post-hoc test (p < 0.01). These findings underscore the potential of azelaic acid as a potent quorum sensing inhibitor, capable of attenuating virulence without exerting direct selective pressure on bacterial growth.

### 3.7 Bioinformatic findings

The described characteristics include the logP, which indicates solubility. As shown in Table 3, azelaic acid has a logP of 1.8863, indicating water solubility. A low logP is associated with greater biological activity. The PSA value indicates the polar surface area of a molecule and how easily it can penetrate the cell. It depends on the type of atoms in the molecule, especially those with higher electronegativity. This value should be less than 140 Å. Azelaic acid has a PSA of 77.57 Å², so the molecule can easily penetrate the cell membrane. The HBD and HBA atoms are useful for determining if the molecule is specific and promotes bioavailability. Molar refractivity (MR), polarizability (P), and polar surface area (PSA) descriptors play an important role in estimating activity and are generally related to drug-active site interactions. On the other hand, a molar refractivity value close to 70 is associated with increased activity. Low polarizability (less than 50) is present in molecules with specific sites of action, meaning they cannot bind to multiple ligands. The data in Table 5 indicate that azelaic acid is a theoretically bioactive compound.

**Table 3.** Molecular and chemoinformatic properties of Azelaic acid.

| **Properties** | **Azelaic acid** |
| --- | --- |
| MF | C<sub>9</sub>H<sub>16</sub>O<sub>4</sub> |
| MW | 188.2209 |
| No. Acceptors | 2 |
| No. Donors | 2 |
| PSA | 77.57 Å<sup>2</sup> |
| LogP | 1.8863 |
| δ | 1.131 ± 0.06 g/cm<sup>3</sup> |
| ST | 46.2 ± 3.0 dyne/cm |
| Polarizability | 18.58 ± 0.5 10<sup>-24</sup>cm<sup>3</sup> |
| MR | 46.87 ± 0.3 cm<sup>3</sup> |
| MV | 166.3 ± 3.0 cm<sup>3</sup> |
| Nrotb | 8 |
| Lipinski's rules Violations | 0 violation |
| SMILES | O=C(O)CCCCCCC(=O)O |

(MF= Molecular formula, MW= Molecular weight (g/mol), HBA= Hydrogen bond acceptors, HBD: Hydrogen bond donors, PSA= Polar Surface Area (Å<sup>2</sup>), LogP= Partition coefficient, δ= density (g/cm<sup>3</sup>), ST= Surface tension, MR= Molar refractivity (cm<sup>3</sup>), nrotb= No. of rotatable bonds, MV= Molar volume (cm<sup>3</sup>), SMILES= Simplified Molecular Input Line Entry System.)

The Osiris Property Explorer and pkCSM simulators were used to predict the bioactivity of azelaic acid. Drug design and the development of new therapeutic alternatives hinge primarily on identifying the toxicity of the bioactive compound. According to predictions, azelaic acid should not exhibit tumorigenic, irritant, reproductive, or mutagenic effects, as shown in Table 4. The main target sites include: nuclear receptors, fatty acid-binding protein family and enzyme inhibitors.

**Table 4.** Prediction of toxicity of azelaic acid.

| **Compound** | **Rep. Ef.** | **Tum.** | **Mut** | **Irr** |
| --- | --- | --- | --- | --- |
| **Azelaic acid** | No | No | No | No |

(Rep. Ef.= Reproductive effective, Tum.= Tumorigenic, Mut.= Mutagenic, Irr= Irritant.)

Analyzing the chemoinformatics data describing its pharmacokinetics, azelaic acid is a promising compound for antimicrobial therapy, theoretically exhibiting good bioavailability and distribution within the body. This is corroborated by the absorption and metabolism data shown in Table 5, making it a good candidate as a bioactive molecule.

**Table 5.** Data of chemoinformatic properties of ADME processes.

| **ADMET** |  | **Azelaic acid** |
| --- | --- | --- |
| **Absorption** | WS | -1.631 |
|  | IA (%) | 92.806 |
|  | SP | -2.732 |
|  | SGP | No |
|  | P-GP-1-In | No |
| **Distribution** | BBBP | -0.388 |
|  | CNSP | -2.945 |
| **Metabolism** | CYP3A4 inhibitor | No |
|  | CYP1A2 inhibitor | No |
| **Excretion** | TC | 1.47 |

(WS: water solubility (log mol/L), IA: intestinal absorption, SP: skin permeability (logKp), SGP: P-glycoprotein substrate, P-GP-1-In: P-glycoprotein I inhibitor, BBBP: blood brain barrier permeability (logBB), CNSP: CNS permeability (logPS), TC: total clearance (log mL/min/kg).)

Azelaic acid has good aqueous solubility, which facilitates its dissolution in biological media. This is compatible with both topical and oral formulations. Intestinal absorption is relatively high, meaning that, in theory, it could be administered orally with moderate bioavailability. A negative SP value indicates low, but not zero, permeability through the skin; that is, although it does not penetrate deeply, it is sufficient to act locally in the epidermis. The SGP value indicates that it is not actively expelled by efflux transporters, which would favor its retention in the tissue where it is applied. Furthermore, since it is not a substrate of P-GP-1-In, it does not interfere with transporters, reducing the risk of drug interactions. Regarding distribution, azelaic acid is not significantly distributed to the brain, which reduces neurological side effects and strengthens its safety profile for topical use. The metabolism is safe, which is consistent with its prolonged use in dermatological treatments. The TC value suggests moderate elimination from the body; if absorbed systemically, the drug can be eliminated relatively efficiently, avoiding accumulation.

### 3.8 Structural and Ionization Analysis of Azelaic Acid

Conformational analysis of azelaic acid predicted a stable, extended zigzag geometry, minimizing steric hindrance along the aliphatic chain (Figure 2). At physiological pH (7.4), the ligand is predicted to exist in dianionic state, with both carboxyl groups fully deprotonated, as shown by the negative charges on the terminal oxygens.

**Figure 2.**
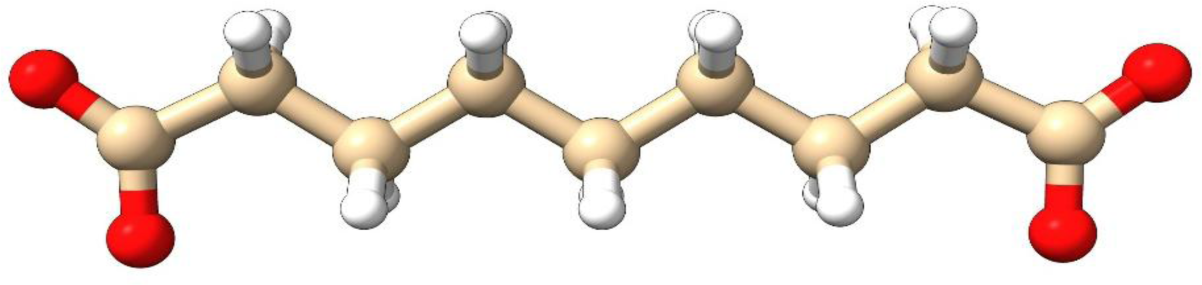
Structure of azelaic acid

### 3.9 Molecular docking

Molecular docking of azelaic acid into LasI (PDB ID: 1RO5) was performed using a binding pocket defined according to previous literature and validated *in silico* with the known LasI inhibitor TZD-C8. Docking of TZD-C8 yielded a binding affinity of −6.9 kcal/mol, forming hydrogen bonds with Arg30 and Ile107, and hydrophobic interactions with Ile56, Trp69, Phe105, Phe117, Thr121, Ala124 and Val148 (Fig 3a), consistent with previously reported inhibitory interactions (42–45). Using the same binding site as a reference, azelaic acid exhibited a calculated binding affinity of −6.1 kcal/mol, establishing hydrogen bonds with Ala106 and Ile107, and hydrophobic contacts with Trp69, Phe105, Phe117, Thr121, Ala124, and Val148 (Fig. 3b and Table 6).

**Figure 3.**
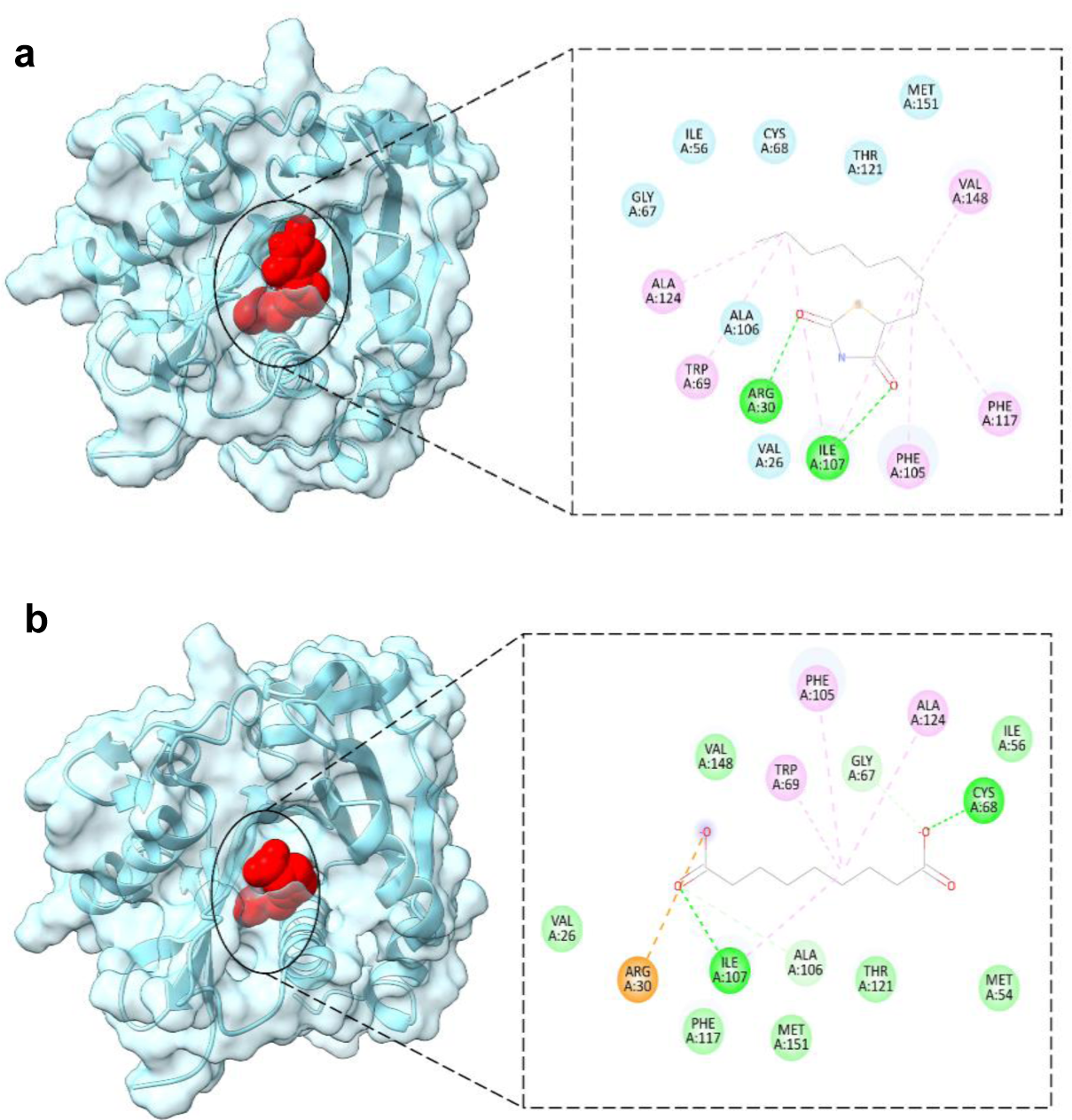
Molecular docking of LasI protein (PDB ID:1RO5). The ligand-binding domain is highlighted in red, and the 2D interactions of the ligands with the receptor are shown: (a) TZD-C8 and (b) azelaic acid.

**Table 6.**
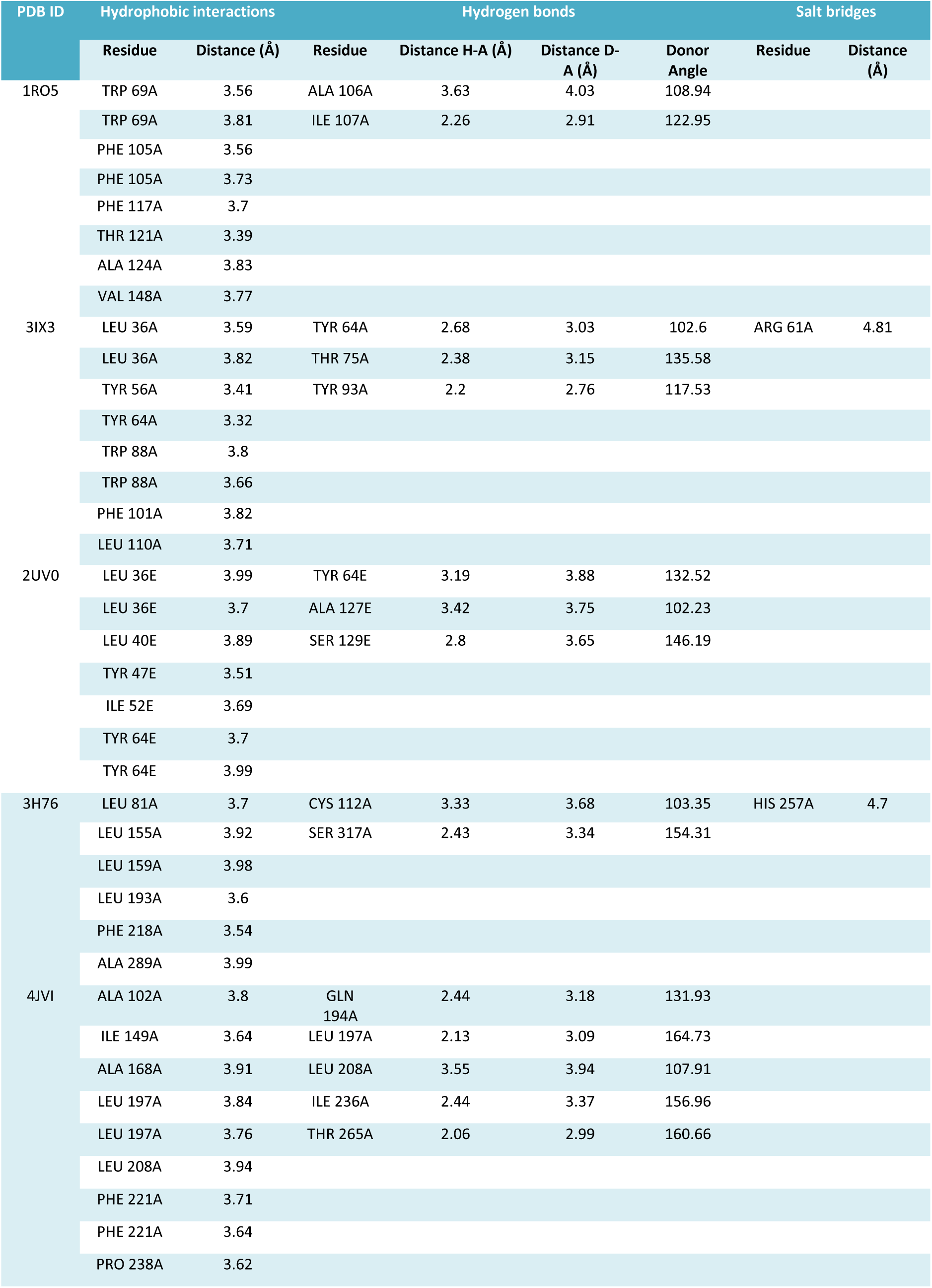
Protein–Ligand Interaction Profiler (PLIP) analysis of azelaic acid binding to selected target proteins.

For the LasR protein (PDB ID: 3IX3 chain A), redocking of its native ligand, OHN (N-3-oxo-dodecanoyl-L-homoserine lactone), yielded a binding affinity of −8.0 kcal/mol and showed hydrogen bond interactions primarily with Asp73 and Ser129. The root mean square deviation (RMSD) of the redocking was 1.459 Å, validating the docking protocol (Fig. 4a). Docking of azelaic acid into LasR (PDB ID: 3IX3) resulted in a predicted binding affinity of −6.5 kcal/mol, maintaining hydrogen bond interactions with Tyr64, Thr75 and Tyr93 and hydrophobic interactions were observed with Leu36, Tyr56, Tyr64, Trp88, Phe101 and Leu110 and a salt bridge with Arg61 (Fig. 4a and Table 6). These residues are consistent with previously reported active-site residues involved in OHN binding, including Tyr56, Trp60, Arg61, Asp73, Thr75, and Ser129, which are conserved across LuxR homologs (46).

**Figure 4.**
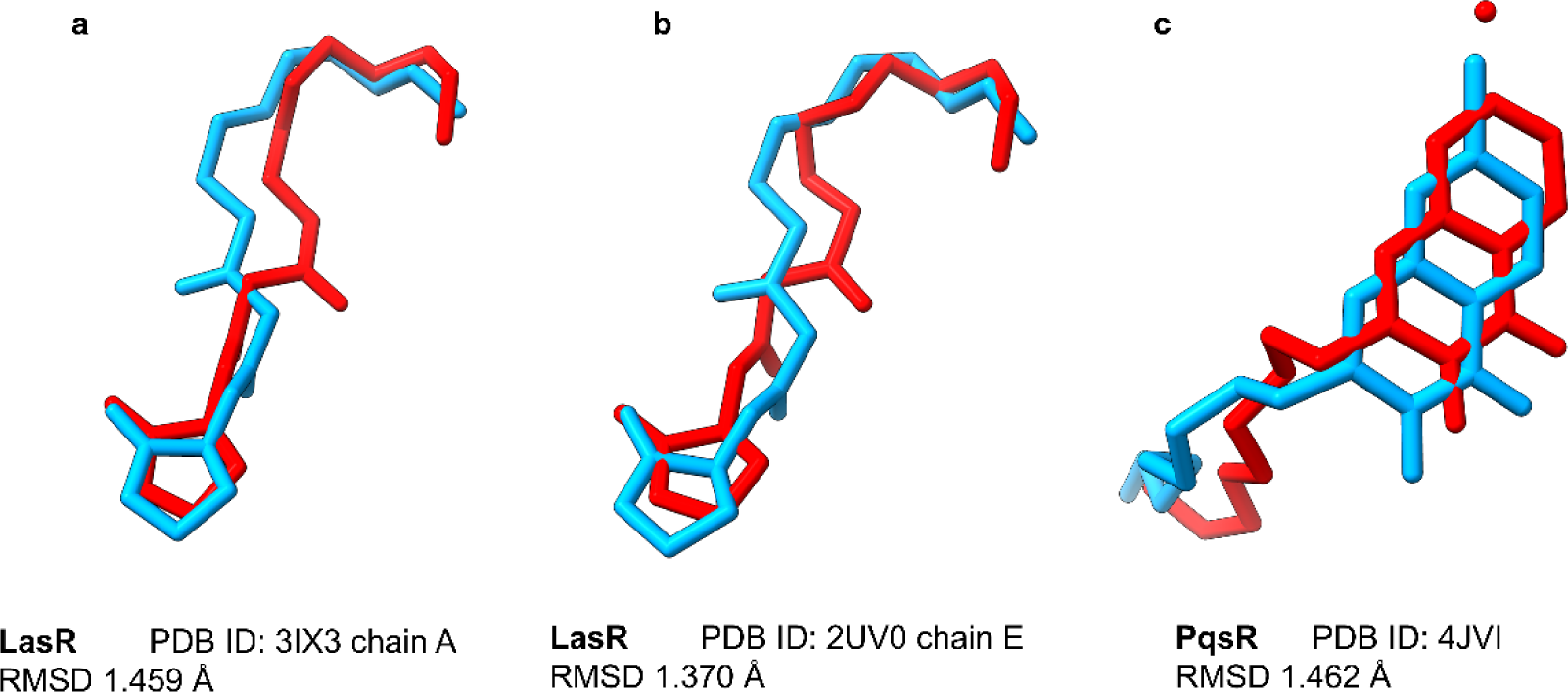
Re-docking analysis. The docking method reproduces the poses of co-crystallized ligands (light blue) with RMSD < 2 Å compared to the re-docked ligands (red) for: (a) LasR (PDB ID: 3IX3), (b) LasR (PDB ID: 2UV0), and (c) PqsD (PDB ID: 4JVI).

For another LasR protein (PDB ID: 2UV0 chain E), redocking of OHN produced a binding affinity of −8.4 kcal/mol and an RMSD of 1.370 Å (Fig 4b). The ligand formed hydrogen bonds with Tyr64, Thr75, and Ser129, as well as additional contacts with Leu40, Tyr47, Val76, Trp88, and Leu125. Azelaic acid docked into 2UV0 showed a predicted binding affinity of −6.4 kcal/mol, forming hydrogen bonds with Tyr64, Ala127 and Ser129 and hydrophobic interactions with Leu36, Leu40, Tyr47, Ile52 and Tyr64 (Fig.5b and Table 6), residues that constitute the active-site pocket (47).

**Figure 5.**
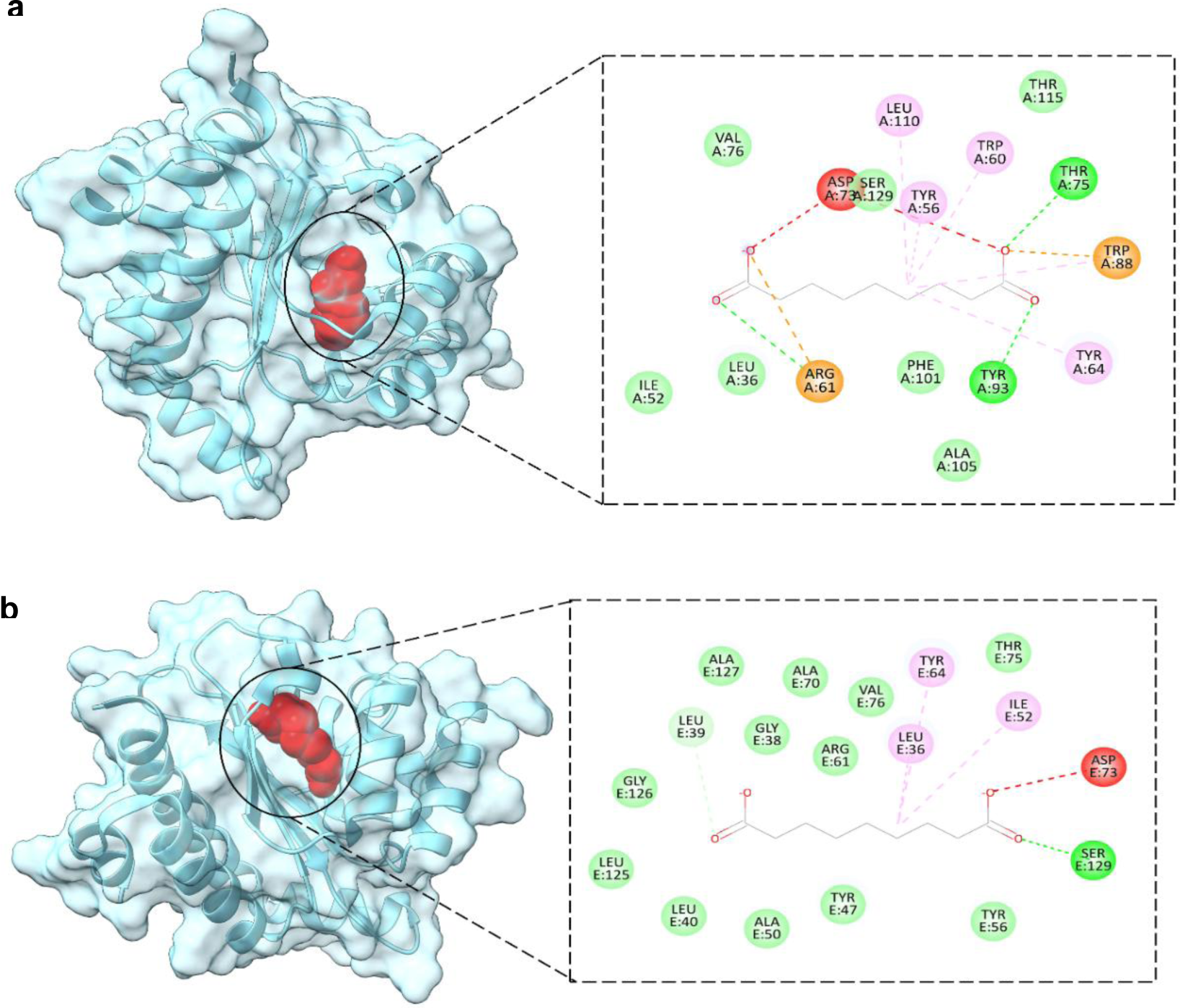
in red and illustrates the 2D interactions of azelaic acid with the protein: (a) 3IX3_chain A and (b) 2UV0_chain E.

Docking with PqsD monomer protein (PDB ID: 3H76) was performed using the amino acids constituting the reported active site to define the docking grid, including Leu81, Cys112, Leu155, Phe218, Met220, Met225, His257, Pro259, and Asn287 (31), with the catalytic triad formed by Cys112, His257, and Asn287 (48). Under these simulated parameters, azelaic acid exhibited a predicted binding affinity of −5.3 kcal/mol, forming hydrogen bonds with Cys112 and Ser317, salt bridge with His257 and hydrophobic interactions with Leu81, Leu155, Leu159, Leu193, Phe218 and Ala289, corresponding to residues of the active pocket (Fig. 6a and Table 6).

**Figure 6.**
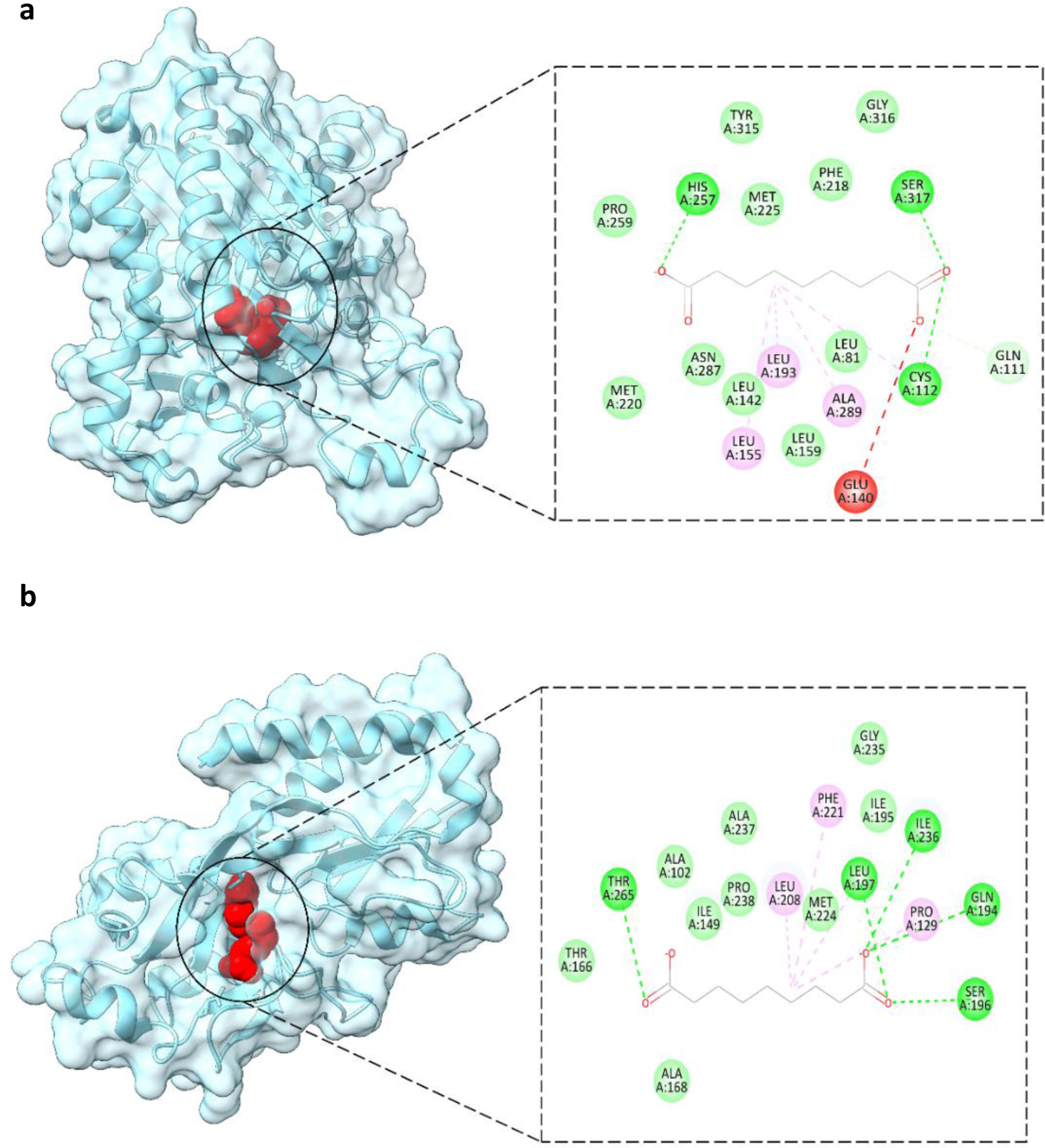
Molecular docking results of azelaic acid with (a) PqsD protein (PDB ID: 3H76) and (b) PqsR protein (PDB ID: 4JVI). The images highlight the ligand-binding domains in red and illustrate the specific 2D interactions of azelaic acid within each catalytic pocket.

For PqsR protein (PDB ID: 4JVI), redocking of QZN (3-amino-7-chloro-2-nonylquinazolin-4(3H)-one) produced a binding affinity of −7.5 kcal/mol and an RMSD of 1.462 Å. The ligand formed contacts with Ala168, Val170, Leu189, Gln194, Leu197, Leu207, Leu208, Arg209, Ile236, Tyr258, Ile263 and Thr265. Azelaic acid docked into 4JVI showed a predicted binding affinity of −6.1 kcal/mol, forming hydrogen bonds with Gln194, Leu197, Leu208, Ile236 and Thr265 and hydrophobic interactions with Ala102, Ile149, Ala168, Leu197, Leu208, Phe221 and Pro238 (Fig. 6b and Table 6), co-localizing with residues within the targeted active-site pocket.

### 3.10 *In Vitro* and *In Vivo* Biocompatibility and Safety Profile of Azelaic Acid

To confirm the biocompatibility and translational safety of azelaic acid (AzA) at the concentrations used for anti-virulence activity, in vitro assays (PBMC viability and erythrocyte hemolysis) were conducted, alongside an *in vivo* toxicity assay using *Tenebrio molitor*.

The *in vitro* evaluation demonstrated an excellent safety profile within the anti-virulence therapeutic range (250 to 750 µg/mL). The MTT assay on human PBMCs revealed that cell viability consistently remained above 90%, indicating an absence of significant cytotoxicity. Concurrently, hemolytic activity against human erythrocytes at these concentrations was strictly below 4% (e.g., 3.73% at 750 µg/mL).

Notably, *in vivo* safety was confirmed via Kaplan-Meier survival curves in the *Tenebrio molitor* model. Larvae injected solely with AzA at the highest experimental concentration (1000 µg/mL) exhibited a 100% survival rate throughout the observation period, matching the results of the PBS-treated control group. This absence of systemic lethality in vivo demonstrates that, even at the minimum inhibitory concentration (MIC), AzA is highly biocompatible and well-tolerated by the host organism, reinforcing its potential as a non-toxic anti-virulence therapy Table 7.

**Table 7.** Cell viability assays using MTT and blood compatibility assays via hemolysis for azelaic acid.

| Compound | Concentration (µg/mL) | Cell Viability (%) | Cell Inhibition (%) | Total Hemolysis (%) |
| --- | --- | --- | --- | --- |
| Azelaic acid | 1000 | 88.250 | 11.750 | - |
|  | 750 | 90.883 | 9.117 | 3.74 |
|  | 500 | 91.267 | 8.733 | 3.59 |
|  | 300 | 93.638 | 6.362 | 3.23 |
|  | 200 | 96.408 | 3.592 | 2.612 |
|  | 100 | 98.108 | 1.893 | 2.56 |
|  | 10 | 98.995 | 1.005 | 2.41 |
|  | 1 | 99.929 | 0.071 | 2.33 |

### 3.11 *In Vivo* Protective Efficacy of Azelaic Acid against *P. aeruginosa* Acute Infection

To determine whether the in vitro attenuation of virulence translated into a protective effect *in vivo*, the survival of *Tenebrio molitor* larvae infected with *P. aeruginosa* (Pa14 and clinical isolate PaBro) was monitored for 72 h and analyzed using Kaplan-Meier survival curves.

The validity of the infection model was confirmed by the PBS-treated control group, which maintained 100% survival throughout the experimental period, indicating that the inoculation procedure itself did not affect larval viability. In contrast, untreated infected larvae showed a marked reduction in survival. Infection with the reference strain (Pa14) resulted in a final survival rate of 0% at 24 h, while the multidrug-resistant clinical isolate (PaBro) produced a comparable mortality level, with a final survival of 0%, confirming the pathogenicity of both strains in this model.

Treatment of both *P. aeruginosa* strains with sub-inhibitory concentrations of azelaic acid (AzA) significantly improved larval survival compared to the corresponding untreated groups (Log-rank test, p < 0.001) Figure 7. For the PaBro isolate, increasing AzA concentrations (250, 500, and 750 µg/mL) was associated with a progressive increase in survival, reaching 70% at the highest concentration. A similar protective effect was observed in larvae infected with the Pa14 strain, indicating that the survival benefit was reproducible in both the reference and clinical isolates.

**Figure 7.**
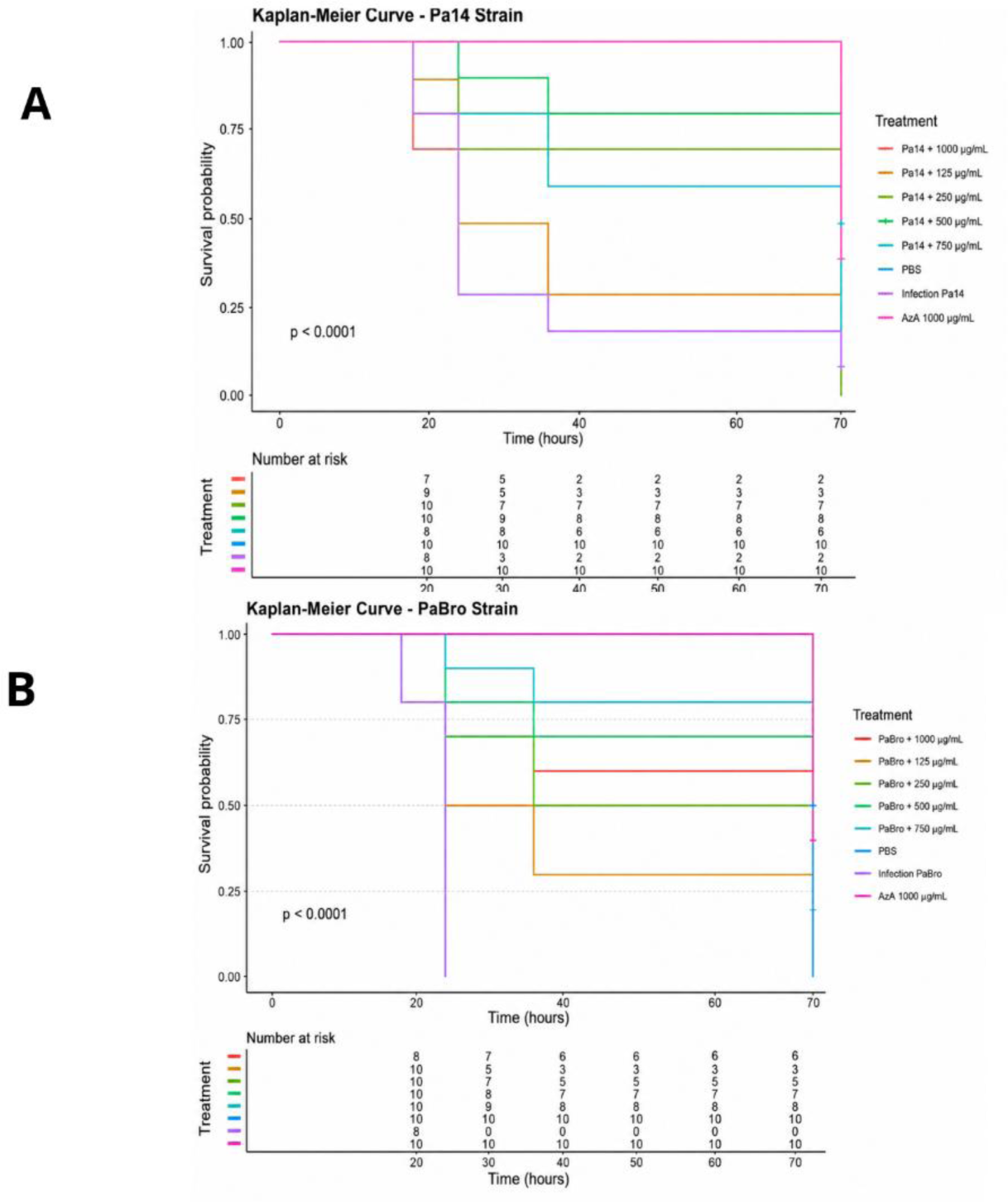
Kaplan-Meier survival curves evaluating the effect of different treatments against bacterial infections. Survival probability over time (hours) is shown for various treatment conditions, controls, and dosages. (A) Survival curve for strain Pa14. (B) Survival curve for strain PaBro. In both graphs, the lines represent the different experimental groups: treatments with varying concentrations of the tested compound combined with the bacteria, a PBS negative control (light blue line), an untreated infection control (purple line), and a toxicity control for compound AzA at 1000 µg/mL without infection (top pink line). The “Number at risk” table corresponding to the evaluated time intervals is detailed at the bottom of each panel. The statistical significance value is indicated in each panel (p < 0.0001, calculated using the log-rank test).

To further evaluate the safety of the compound in vivo, healthy larvae were injected with AzA (1000 µg/mL) in the absence of bacterial infection. This group maintained 100% survival during the 72-hour observation period, comparable to that of the PBS-treated control, indicating that AzA did not produce detectable systemic toxicity at the highest concentration evaluated Figure 7.

Overall, these findings demonstrate that AzA significantly improves host survival during *P. aeruginosa* infection without exhibiting detectable toxicity under the experimental conditions, supporting its potential as an anti-virulence therapeutic candidate.

### 3.12 Effect of azelaic acid on pyocyanin production *in vivo*

To determine whether the attenuation of virulence observed in *in vitro* assays was replicated in a biological system, pyocyanin production was quantified directly from *Tenebrio molitor* larvae infected with *P. aeruginosa* strains Pa14 and PaBro. Consistent with results obtained in liquid culture, treatment with azelaic acid (AzA) significantly reduced pigment accumulation in the hemocoel and larval tissues in a dose-dependent manner. High concentrations of the characteristic pigment were evident in the infected, untreated groups, whereas a drastic reduction in recovered pyocyanin levels was observed in larvae treated with sub-MIC concentrations of AzA (250, 500, and 750 µg/mL). For Pa14, reductions of 40%, 71%, and 90% were observed at 250, 500, and 750 µg/mL, respectively; for PaBro, the reduction ranged from 29% at 250 µg/mL to 73% at 750 µg/mL. This protective metabolic effect correlated with a significant increase in larval survival rates—analyzed via Kaplan-Meier curves—demonstrating that AzA can mitigate the expression of this key virulence factor during active infection without exhibiting intrinsic toxicity toward the animal model at concentrations up to 1000 µg/mL Figure 8.

**Figure 8.**
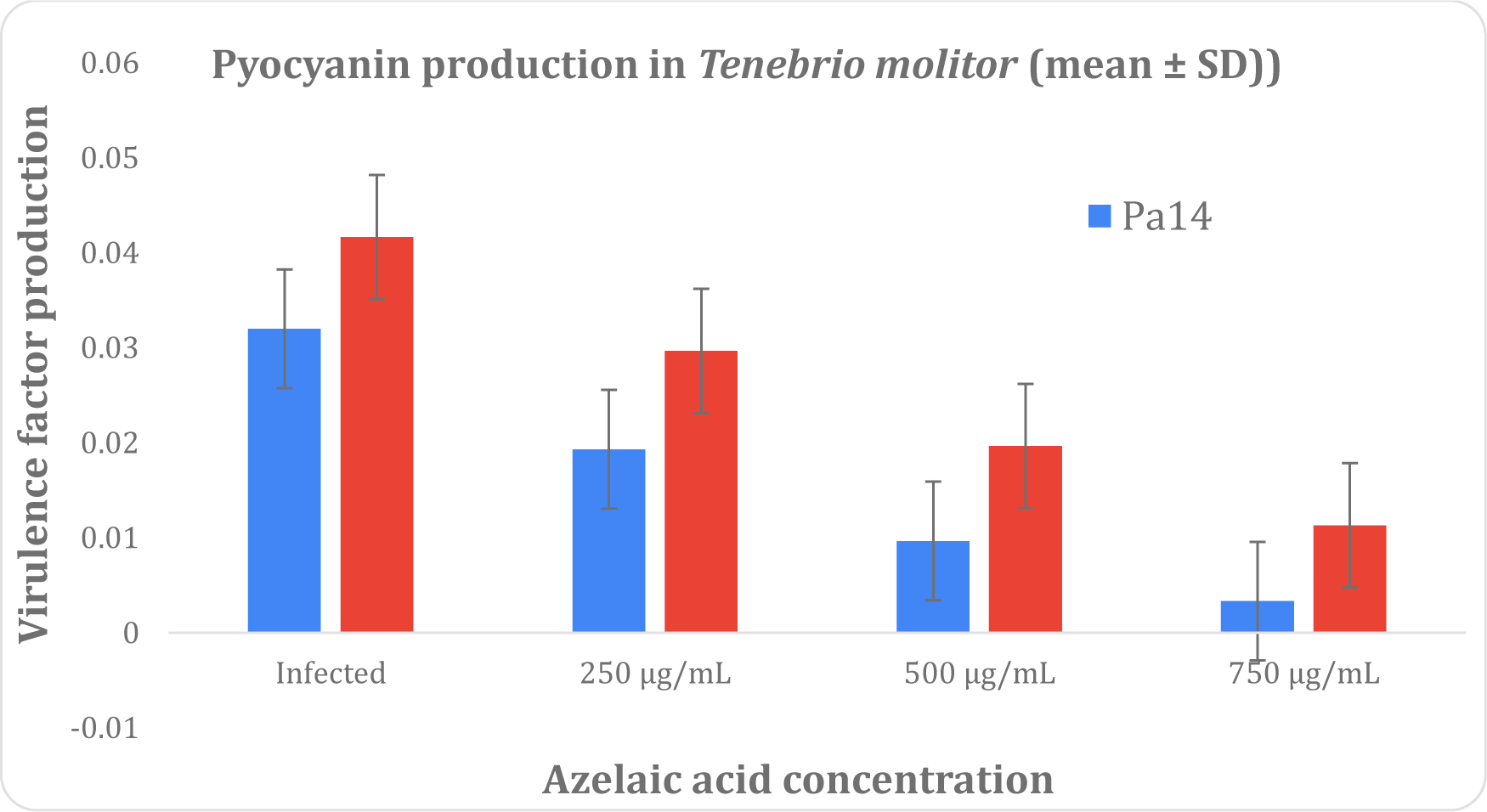
Effect of azelaic acid (AzA) on *in vivo* pyocyanin production in the *Tenebrio molitor* infection model. Larvae were infected with *Pseudomonas aeruginosa* strains Pa14 and PaBro and treated with sub-inhibitory concentrations of AzA (250, 500, and 750 µg/mL. Spectrophotometric quantification of pyocyanin extracted via mechanical lysis and pH-gradient purification. Data represent the mean standard deviation (SD) of three independent experiments.

## Discussion

The results of this study demonstrate that azelaic acid acts as an agent with a significant capacity to interfere—in a dose-dependent manner—with the virulence mechanisms of *Pseudomonas aeruginosa* coordinated by bacterial intercellular communication (quorum sensing or QS). Notably, the inhibition of critical factors such as pyocyanin, alginate, elastase, and protease was consistently observed across both reference strains and clinical isolates exhibiting multidrug-resistant (MDR) and pandrug-resistant (PDR) profiles. Beyond its *in vitro* effects, it also possesses the ability to reduce factors like pyocyanin *in vivo*, thereby increasing survival rates in infected models such as *Tenebrio molitor*.

Regarding its bactericidal capacity, while mild-to-moderate activity has previously been reported, this study tested its ability to inhibit the growth of various Gram-negative and Gram-positive strains—including both reference strains and clinical isolates—yielding MIC values between 250 µg/mL and 1000 µg/mL. Although these values may appear high, they are actually low when compared to figures previously reported by other research groups. This finding is particularly significant, as it suggests that azelaic acid may interfere with regulatory metabolic pathways independent of traditional antibiotic resistance mechanisms regarding its antibacterial activity.

As an antibacterial agent, AzA exhibits activity against bacteria such as *Cutibacterium acnes* (formerly *Propionibacterium acnes), Staphylococcus epidermidis, Staphylococcus aureus,* and *Pseudomonas aeruginosa*. Its efficacy is dependent on concentration and pH, with more pronounced effects observed at low pH levels and high concentrations (49). While primarily used in dermatology to treat various skin conditions, in the context of infections, it is currently employed to treat several types of acne, as well as rosacea and acne vulgaris (50). Its use in dermatology attests to the safety of AzA and positions it as a promising molecule for treating infections caused by bacteria classified under the ESKAPE acronym. This was further confirmed by safety studies involving PBMC cytotoxicity and hemolysis.

Blaskovich(51) demonstrated MICs for 6 strains of *S. aureus* ATCC (two MRSA, one MSSA, one GISA, and one VRSA) ranging from 8000 to 16000 µg/mL, concentrations much higher than those we reported for the same strain MRSA ATCC 43300, MSSA ATCC 6538, and a clinical strain *S. aureus* Ur, with MICs of 1000 µg/mL and 250 µg/mL, respectively. For *E. coli* ATCC 25922, they achieved an MIC of 16000 µg/mL, while our results are 1000 µg/mL for said ATCC and 250 µg/mL for the clinical strain Ur.

Charnock (52) showed inhibition of AzA with MICs of 1.5 to 4.5 mg/mL for all *S. aureus* strains using the disk diffusion method. Esterified AzA was also evaluated with MICs of 0.25 to 0.75 mg/mL, considerably better than AzA. The broth microdilution evaluation we performed for the *clinical S. aureus* strain coincided with 0.25 mg/mL; however, for ATCC strains, the MICs were lower than those obtained with AzA in its dicarboxylic state. This may be due to the method used and the diffusion of AzA in the Kirby-Bauer method.

A further study conducted by Muhammet (53) evaluated an AzA gel, achieving up to 20 mm of inhibition for *S. aureus* and up to 10 mm for *E. coli* using the diffusion method; however, the concentration used was not mentioned.

Bandzerewicz et al. (54) used *Pseudomonas aeruginosa* ATCC 9027 and *S. aureus* ATCC 6538 to evaluate the antibacterial activity of an azelaic acid-based biomaterial, demonstrating a significant reduction in growth and viable cells after treatment. This coincides with the results presented in our work against the *Pseudomonas* and *S. aureus* strains. However, the concentrations of Aza used in the study were not mentioned.

The precise mechanisms by which azelaic acid (AzA) exhibits antibacterial activity remain to be fully elucidated, as multiple pathways have been described in the literature. However, effects on key enzymatic activities, macromolecular synthesis, and intracellular pH homeostasis have been strongly suggested (52).

The main mechanism involves the inhibition of bacterial mitochondrial oxidoreductase enzymes, which are essential for microbial respiration and protein synthesis. By acting on this enzyme system, AzA disrupts energy production within bacterial cells, reducing proliferation and ultimately leading to bacterial death. The mechanism of AzA involves the inhibition of bacterial thioredoxin reductase (55), thereby suppressing protein and DNA synthesis (49).

Another reported mechanism for AzA is the reduction of intracellular pH, which affects the maintenance of a pH gradient across the membrane and leads to a loss of energy generation by bacterial metabolism, such as the respiratory chain (where reduced NADH dehydrogenase, succinic acid dehydrogenase, and reduced cytochrome-c ubiquinone oxidoreductase are inhibited) and anaerobic glycolysis (which inhibits hexokinase). This low-energy environment decreases protein synthesis and the synthesis of RNA and DNA (56).

Beyond direct metabolic impairment, communicating microorganisms utilize quorum sensing (QS) to coordinate their gene expression profile, that is, the regulation of virulence factors, biofilm development, secondary metabolite production, and host interaction. Therefore, the search for quorum sensing inhibitors (QSIs) emerges as a new therapeutic approach for bacterial infections (57). Repurposing clinically already used molecules aims to expedite the transition to clinical trials by omitting preliminary safety, pharmacokinetics, and toxicology (LADMET) studies, as these have already been performed.

*Pseudomonas aeruginosa* regulates virulence, biofilm formation, and adaptive responses through interconnected quorum sensing (QS) systems, primarily the Las and PQS regulatory networks (58,59). In the present study, the phenotypic antivirulence effects observed experimentally *in vitro* were complemented by molecular docking analyses to explore potential structural interactions between AzA and key QS-associated proteins. Our *in vitro* assays demonstrated that AzA modulated QS-related phenotypes, reducing the production of key virulence factors and supporting its role as a potential architectural modulator of bacterial communication.

Similar behavioral trends have been reported for other carboxylic derivatives and related compounds. For instance, Liu, et al (60), evaluated a fungal dicarboxylic derivative, the active compound PT22 (1H-pyrrole-2,5-dicarboxylic acid) isolated from *Perenniporia tephropora* FF2, an endophytic fungus of *Areca catechu L*., exhibits QS inhibitory activity against *P. aeruginosa*, where at concentrations (0.50 mg/mL, 0.75 mg/mL and 1.00 mg/mL), it reduces the production of QS-related virulence factors, such as pyocyanin and rhamnolipid, and inhibits the formation of *P. aeruginosa* Pa01 biofilms, despite the structural difference, the behavior is similar to that of azelaic acid. Furthermore, Jabłońska-Wawrzycka (61) demonstrated that coordination complexes with Mn and a carboxylic ligand, such as Mn-pyCOOH-H₂O and [Mn-pyCOOH-H₂O], significantly reduced biofilm growth of the *P. aeruginosa* Pa01 strain. Biofilm formation was inhibited by 48% (Mn-pyCOOH-H₂O complex) and 45% ([Mn-pyCOOH-H₂O]ₙ complex) compared to an untreated control when the concentration of the complexes was 0.5 mM, a concentration similar to that used in our experiments, and that they inhibited alginate by similar percentages.

While the search for novel QS modulators has explored diverse chemical spaces, ranging from natural products like curcumin (62) and indoles (63) to furanones (64), plant extracts (24), and repurposed drugs such as sitagliptin (65), our study positions azelaic acid as an attractive candidate for further evaluation. Unlike its traditional topical application in dermatology, our findings highlight its notable anti-virulence activity in both reference strains and clinical isolates with resistant phenotypes.

Chemoinformatic analysis suggest that AzA aligns with drug-likeness criteria and exhibits a favorable theoretical safety profile, biological validation remains crucial.

To complement the *in vitro* findings, molecular docking simulations were utilized to explore the structural feasibility of azelaic acid interacting with key quorum sensing targets. It is important to emphasize that these computational models are intended to serve as supportive structural evidence rather than a definitive mechanistic demonstration. Azelaic acid exhibited moderate predicted binding affinity (−6.1 kcal/mol), toward LasI (PDB ID: 1RO5), interacting within the catalytic cavity and establishing contacts with residues such as Ile107, which have been previously implicated in ligand stabilization (30). Although the predicted affinity was lower than that of optimized synthetic inhibitors, its positioning within the active site suggests a possible competitive accommodation within the catalytic pocket that may contribute to reduced AHL production.

For LasR (3IX3 and 2UV0), azelaic acid maintained interactions within the conserved ligand-binding domain, particularly involving Thr75 and Ser129. While its predicted binding energy was lower than that of the native autoinducer OHN (−8.0 to −8.4 kcal/mol), the compound adopted a comparable binding orientation within the conserved pocket. Several previously reported LasR inhibitors interact with residues such as Tyr56, Asp73, Thr75, and Ser129 (66, 67, 68) and the interaction profile observed here aligns with these established binding patterns. These findings suggest that azelaic acid may exert a modulatory effect on LasR-mediated transcriptional activation rather than acting as a high-affinity competitive inhibitor.

Regarding the PQS pathway, docking simulations suggest that azelaic acid could accommodate within the active site region of PqsD (PDB ID: 3H76). At physiological pH (7.4), the dianionic form of the ligand is predicted to form a salt bridge with His257 and hydrogen bonds with Cys112, residues that constitute the Cys112-His257-Asn287 catalytic triad. The engagement of Cys112 is particularly significant, as this residue is responsible for the initial nucleophilic attack during anthraniloyl-CoA processing (31, 38). While the predicted binding affinity (−5.3 kcal/mol) indicates a modest interaction, the orientation of the carboxylate groups toward these key residues could potentially interfere with substrate binding or catalysis.

Docking against PqsR (PDB ID: 4JVI) further revealed interactions with key binding-pocket residues, including Gln194 and Ile236, consistent with regions previously implicated in ligand stabilization (69). Previous structural studies have reported that synthetic inhibitors form hydrogen bonds with residues such as Gln194 and Arg209 while occupying a hydrophobic cavity defined by Ile236–Pro238 and Phe221–Met224. Although azelaic acid exhibited a moderate predicted binding affinity (−6.1 kcal/mol), its positioning within this conserved pocket aligns with these characterized binding modes, supporting a potential modulatory interaction with the PQS signaling pathway.

Importantly, although docking energies did not indicate high-affinity inhibition when compared to native ligands or optimized QS inhibitors, azelaic acid consistently occupied conserved ligand-binding domains across multiple QS-related targets. This broad interaction profile across different receptors offers a plausible structural hypothesis for a cumulative disruption of regulatory pathways involved in bacterial communication and pathogenicity observed *in vitro*. Rather than acting as a potent single-target inhibitor, azelaic acid may exert minor, cumulative modulatory effects across interconnected pathways, ultimately contributing to the overall attenuation of virulence-associated phenotypes.

As previously shown, azelaic acid at sub-inhibitory concentrations reduces four key virulence factors in Pseudomonas aeruginosa. These sub-MIC levels were defined for each of the five strains included in this study, reflecting the operational principle of quorum-sensing inhibitors: doses that do not suppress growth but selectively attenuate bacterial communication and virulence(70).

Taken together, these data support a model in which azelaic acid acts primarily as an anti-virulence modulator rather than as a classic bactericidal agent. Sub-inhibitory (sub-MIC) concentrations were sufficient to attenuate key determinants of *P. aeruginosa* pathogenicity across diverse genetic backgrounds, including multidrug-resistant (MDR) and pandrug-resistant (PDR) clinical isolates. Variations in inhibition stem from phenotypic and genotypic differences among strains, as the pathogenicity islands present in each strain dictate varying degrees of susceptibility; for instance, strain Pa14 is more virulent than Pa01 or the ATCC 27853 reference strain(25). Furthermore, the clinical strains tested exhibited similar behavior, with greater inhibitory activity observed against the PDR strain PaBro compared to the MDR strain PaHer. Clinical strains exhibit significantly higher and more unpredictable resistance to quorum sensing inhibitors (QSIs) than laboratory reference strains. While reference strains (such as *Pseudomonas aeruginosa* Pa01) possess stable genomes and homogeneous communication systems, clinical isolates obtained from actual patients carry pre-existing mutations resulting from the stress of the hospital environment and antibiotic use—as evidenced by the results, where Pa01, Pa14, and PaBro showed greater inhibition compared to PaHer (71,72).

The *in vivo* protection observed in *Tenebrio molitor* further indicates that this phenotypic attenuation translates into reduced pathogenicity in a living host. Pa14 was used as the most virulent type strain, and PaBro for its PDR (pandrug-resistant) profile, to determine whether the treatment was as effective *in vivo* as *in vitro* against both strains. Kaplan-Meier (KM) plots demonstrate the advantage offered by the treatment—even at the lowest concentration tested— compared to untreated, infected controls; concentrations of 500 and 750 µg/mL provided the best protection for the larvae. While this protection may stem from AzA’s antibacterial mechanisms(49–51), *in vivo* pyocyanin quantification was used to verify whether virulence attenuation contributed to larval survival; a reduction of over 70% was observed in treated larvae compared to those infected with Pa14 and PaBro. Pyocyanin is a key redox-active toxin essential for the pathogenicity of *Pseudomonas aeruginosa*. Due to its zwitterionic chemical properties, pyocyanin readily penetrates host cell membranes. Once inside, it reacts directly with cellular oxygen to generate massive amounts of reactive oxygen species (ROS), such as the superoxide anion (O_2_-) and hydrogen peroxide (H_2_O_2_), causing severe oxidative stress and tissue destruction. Blocking pyocyanin eliminates the primary source of free radicals, thereby mitigating cellular necrosis and halting damage to the infected organism’s biological tissues (73,74,75). By inhibiting pyocyanin with azelaic acid, the bacterial strain loses its main weapon for destroying the insect or animal model’s tissues; although the bacteria remain alive within the organism, their lethality is completely neutralized. The suitability of *Tenebrio molitor* as a robust model for *in vivo* antimicrobial screening has been widely supported, demonstrating that survival profiles obtained via Kaplan-Meier curves are comparable to those of traditional mammalian models (76).

Regarding the safety of AzA, and based on international standards ASTM F756 and ISO 10993-5, safety criteria include a hemolysis rate of 0% to 5% and PBMC cell viability of 70% or higher (77,78,79). Our results show 3.74% hemolysis at 750 µg/mL and 90.88% compatibility at the same concentration; notably, compatibility remains at 88.25% at 1000 µg/mL—a finding confirmed by the in vivo model in which this concentration was administered to assess safety. Therefore, the sub-MIC concentrations used are safe for translation to in vivo models. Studies indicate that compound biocompatibility is validated against these thresholds to ensure safety (80).

Taken together, these findings support a model in which azelaic acid acts primarily as an anti-virulence modulator rather than as a classical bactericidal agent. Across reference strains and clinical MDR/PDR isolates, sub-inhibitory concentrations were sufficient to attenuate key pathogenic traits, including pyocyanin, alginate, elastase, and protease production, and these phenotypic effects were mirrored in vivo by improved survival in the *Tenebrio molitor* infection model.

Although the predicted binding affinities were modest, the consistent occupation of conserved ligand-binding sites across multiple quorum-sensing targets provides a plausible structural basis for the cumulative disruption of regulatory pathways. Nevertheless, these docking results should be interpreted as supportive evidence rather than definitive mechanistic proof, and further studies integrating genomics, proteomics, and direct quorum-sensing reporter assays will be necessary to clarify the molecular events underlying the observed phenotypes. In addition, while *T. molitor* is a valuable and reproducible in vivo platform for antimicrobial screening, validation in mammalian infection models will be important to strengthen translational relevance.

In a clinical context, azelaic acid emerges as a promising candidate for repurposing as an anti-virulence adjuvant, particularly in settings where attenuation of virulence and biofilm-associated tolerance may reduce the effective antibiotic dose and potentially delay resistance emergence. Its established use in dermatology, together with the favorable safety profile demonstrated here by PBMC cytotoxicity and hemolysis assays, further supports its potential translational value. In summary, this study positions azelaic acid as a quorum-sensing and virulence modulator in *P. aeruginosa* with consistent in vitro and in vivo activity, making it an attractive candidate for future anti-virulence and adjuvant strategies against resistant infections.

## Conclusion

Azelaic acid demonstrated anti-virulence activity by significantly inhibiting—in a dose-dependent manner—virulence factors regulated by the quorum sensing system in *Pseudomonas aeruginosa,* including multidrug-resistant (MDR) and pandrug-resistant (PDR) clinical isolates. Notably, the marked suppression of pyocyanin, alginate, elastase, and protease production at sub-inhibitory concentrations indicates that its primary activity is associated with virulence attenuation rather than direct bactericidal action.

Complementary molecular docking analyses provided structural support for these findings, suggesting that azelaic acid can occupy conserved binding regions across multiple quorum sensing-related targets, including LasI, LasR, PqsD, and PqsR. Although these simulations do not constitute definitive mechanistic proof, they offer a plausible structural basis for the cumulative disruption of bacterial communication and the pathogenicity observed *in vitro*.

It is important to note that this study provides the first evidence that azelaic acid acts as a quorum sensing inhibitor in *P. aeruginosa*, both *in vitro* and *in vivo*. Given its established safety profile and its ability to attenuate virulence without exerting strong selective pressure, azelaic acid emerges as a promising candidate for repurposing as an anti-virulence agent in the context of antimicrobial resistance. Further mechanistic and *in vivo* validation studies will be essential to determine its therapeutic potential.

## Acknowledgment

The authors express their gratitude to Dr. Lidia Gonzalez Morales and Cristian Sadalis for their technical support, and to the administration and laboratories of the QFB degree program for the support provided in carrying out this work.

## References

1 Ahmed SK, Mohammad NS, Ahmed KA, Ibrahim MS, Hussein SH, Hussein NR, et al. Antimicrobial resistance: impacts, challenges, and future prospects. J Med Surg Public Health. 2024;2:100081. doi: 10.1016/j.glmedi.2024.100081.

2 Pariente N, PLOS Biology Staff Editors. The antimicrobial resistance crisis needs action now. PLoS Biol. 2022;20(11):e3001918. doi: 10.1371/journal.pbio.3001918.

3 Li W, Sun S, Sun Y, Wu X, Shi L, Wang Y, et al. Association between antibiotic resistance and increasing ambient temperature in China: an ecological study with nationwide panel data. Lancet Reg Health West Pac. 2023;30:100628. doi: 10.1016/j.lanwpc.2022.100628.

4 Macesic N, Ginn AM, Harris PNA, Paterson DL, Holmes AH, Llewelyn MJ, et al. Multidrug-resistant Gram-negative bacterial infections. Lancet. 2025;405(10474):257–272. doi: 10.1016/S0140-6736(24)02081-6.

5 Marino A, Maniaci A, Lentini M, Ronsivalle S, Nunnari G, Cocuzza S, et al. The global burden of multidrug-resistant bacteria. Epidemiologia. 2025;6(2):21. doi: 10.3390/epidemiologia6020021.

6 Spagnolo AM, Sartini M, Cristina ML. Pseudomonas aeruginosa in the healthcare facility setting. Rev Med Microbiol. 2021;32(3):169–175. doi: 10.1097/MRM.0000000000000271.

7 Moradali MF, Ghods S, Rehm BHA. Pseudomonas aeruginosa lifestyle: a paradigm for adaptation, survival, and persistence. Front Cell Infect Microbiol. 2017;7:39. doi: 10.3389/fcimb.2017.00039.

8 Reynolds D, Kollef M. The epidemiology, pathogenesis, and treatment of Pseudomonas aeruginosa infections: an update. Drugs. 2021;81(18):2117–2131. doi: 10.1007/s40265-021-01635-6.

9 Qin S, Xiao W, Zhou C, Pu Q, Deng X, Lan L, et al. Pseudomonas aeruginosa: pathogenesis, virulence factors, antibiotic resistance, interaction with host, technology advances and emerging therapeutics. Signal Transduct Target Ther. 2022;7:199. doi: 10.1038/s41392-022-01056-1.

10 Miranda SW, Asfahl KL, Dandekar AA, Greenberg EP. Pseudomonas aeruginosa quorum sensing. Adv Exp Med Biol. 2022;1386:95–115. doi: 10.1007/978-3-031-08491-1_4.

11 Raya J, Montagut EJ, Marco MP. Analysing the integrated quorum sensing (IQS) system and its potential role in Pseudomonas aeruginosa pathogenesis. Front Cell Infect Microbiol. 2025;15:1575421. doi: 10.3389/fcimb.2025.1575421.

12 Long Y, Li Z, Li M, Lu P, Deng Y, Wu P, et al. Pseudomonas aeruginosa PQS quorum sensing mediates interaction with Mycobacterium abscessus in vitro. Microorganisms. 2025;13:116. doi: 10.3390/microorganisms13010116.

13 Gou L, et al. Revealing quorum-sensing networks in Pseudomonas aeruginosa infections through internal and external signals to prevent new resistance trends. Microbiol Res. 2024;289:127915. doi: 10.1016/j.micres.2024.127915.

14 Lin J, et al. Quorum sensing in Pseudomonas aeruginosa and its relationship to biofilm development. Methods Enzymol. 2019;1323:1–16. doi: 10.1021/bk-2019-1323.ch001.

15 de Kievit TR, Iglewski BH. Role of the Pseudomonas aeruginosa las and rhl quorum-sensing systems in rhlI regulation. FEMS Microbiol Lett. 2002;212(1):101–106. doi: 10.1016/S0378-1097(02)00735-8.

16 Bru JL, Rawson B, Trinh C, Whiteson K, Høyland-Kroghsbo NM, Siryaporn A. PQS produced by the Pseudomonas aeruginosa stress response repels swarms away from bacteriophage and antibiotics. J Bacteriol. 2019;201:e00383–19. doi: 10.1128/JB.00383-19.

17 Fleitas Martínez O, Cardoso MH, Ribeiro SM, Franco OL. Recent advances in anti-virulence therapeutic strategies with a focus on dismantling bacterial membrane microdomains, toxin neutralization, quorum-sensing interference and biofilm inhibition. Front Cell Infect Microbiol. 2019;9:74. doi: 10.3389/fcimb.2019.00074.

18 Ma ZP, Song Y, Cai ZH, Lin ZJ, Lin GH, Wang Y, et al. Anti-quorum sensing activities of selected coral symbiotic bacterial extracts from the South China Sea. Front Cell Infect Microbiol. 2018;8:144. doi: 10.3389/fcimb.2018.00144.

19 Omwenga EO, Hensel A, Pereira S, Shitandi AA, Goycoolea FM. Antiquorum sensing, antibiofilm formation and cytotoxicity activity of commonly used medicinal plants by inhabitants of Borabu sub-county, Nyamira County, Kenya. PLoS One. 2017;12(11):e0185722. doi: 10.1371/journal.pone.0185722.

20 Zhao YM, Zhang QY, Zhang L, Bao YL, Guo YT, Huang LR, et al. Inhibition of quorum sensing-mediated biofilm formation and spoilage factors in Pseudomonas fluorescens by plasma-activated water. Foods. 2025;14:3773. doi: 10.3390/foods14213773.

21 Karpiñski TM, Adamczak A. Fucoxanthin—an antibacterial carotenoid. Antioxidants (Basel). 2019;8(8):239. doi: 10.3390/antiox8080239.

22 Clinical and Laboratory Standards Institute (CLSI). Methods for dilution antimicrobial susceptibility tests for bacteria that grow aerobically; approved standard—eighth edition. Wayne (PA): CLSI; 2009.

23 Adamczak A, Oarowski M, Karpiñski TM. Curcumin, a natural antimicrobial agent with strain-specific activity. Pharmaceuticals (Basel). 2020;13(7):153. doi: 10.3390/ph13070153.

24 Lugo-Gutiérrez DA, Martínez-González G, Santos-López CS, Almeida-Villegas JA. First report: Tagetes erecta inhibits quorum-sensing in Pseudomonas aeruginosa. Ars Pharm. 2026;67(2):188–199. doi: 10.30827/ars.v67i2.35452.

25 Martínez-González G, et al. Molecular Insight into the Inhibition of Pseudomonas aeruginosa Quorum Sensing by 5-Bromoindole-3-carboxaldehyde and Curcumin: A Combined In Vitro Study, Molecular Dynamics, and Docking Studies. ACS Omega. 2026. doi: 10.1021/acsomega.6c02161.

26 Knutson CA, Jeanes A. A new modification of the carbazole analysis: application to heteropolysaccharides. Anal Biochem. 1968;24(3):470–481. doi: 10.1016/0003-2697(68)90154-1.

27 Ohman DE, Cryz SJ, Iglewski BH. Isolation and characterization of Pseudomonas aeruginosa PAO1 mutant that produces altered elastase. J Bacteriol. 1980;142(3):836–842. doi: 10.1128/jb.142.3.836-842.1980.

28 Alisaac A. In silico analysis of quorum sensing modulators: insights into molecular docking and dynamics and potential therapeutic applications. PLoS One. 2025;20(6):e0325830. doi: 10.1371/journal.pone.0325830.

29 Gould TA, Schwelzer HP, Churchill MEA. Structure of the Pseudomonas aeruginosa acyl-homoserine lactone synthase LasI. Mol Microbiol. 2004;53(4):1135–1146. doi: 10.1111/j.1365-2958.2004.04211.x.

30 Singothu S, Begum PJ, Maddi D, Devsani N, Bhandari V. Unveiling the potential of marine compounds as quorum sensing inhibitors targeting Pseudomonas aeruginosa LasI: a computational study using molecular docking and molecular dynamics. J Cell Biochem. 2023;124(10):1573–1586. doi: 10.1002/jcb.30465.

31 Martin KDG, Padilla KGV, Buan IJA. In silico study of Ayapana triplinervis bioactive compounds against quorum-sensing system of Pseudomonas aeruginosa. Orient J Chem. 2021;37(1):143–150. doi: 10.13005/ojc/370119.

32 Schake P, Bolz SN, Linnemann K, Schroeder M. PLIP 2025: introducing protein-protein interactions to the protein-ligand interaction profiler. Nucleic Acids Res. 2025;53(W1):W463–W465. doi: 10.1093/nar/gkaf361.

33 Li DD, Wang Y, Li H, Niu WX, Hong J, Jung JH, et al. Multifaceted Antipathogenic Activity of Two Novel Natural Products, Chermesiterpenoid B and Chermesiterpenoid B Seco Acid Methyl Ester, Against Pseudomonas aeruginosa. Microb Biotechnol. 2025;18(2):e70101. doi: 10.1111/1751-7915.70101.

34 Petronio Petronio G, Pietrangelo L, Cutuli MA, Magnifico I, Venditti N, Guarnieri A, et al. Emerging Evidence on Tenebrio molitor Immunity: A Focus on Gene Expression Involved in Microbial Infection for Host-Pathogen Interaction Studies. Microorganisms. 2022;10(10):1983. doi: 10.3390/microorganisms10101983.

35 Moret Y. “Trans-generational immune priming”: specific enhancement of the antimicrobial immune response in the mealworm beetle, Tenebrio molitor. Proc Biol Sci. 2006;273(1592):1399–1405. doi: 10.1098/rspb.2006.3465.

36 Ham S, Kim H, Jo MJ, Lee J, Byun Y, Ko G, et al. Combined Treatment of 6-Gingerol Analog and Tobramycin for Inhibiting Pseudomonas aeruginosa Infections. Microbiol Spectr. 2021;9(2):e00192–21. doi: 10.1128/Spectrum.00192-21.

37 Stalpers LJA, Kaplan EL. Edward L. Kaplan and the Kaplan-Meier Survival Curve. BSHM Bull. 2018;33(2):109–135. doi: 10.1080/17498430.2018.1450055.

38 Amin K, Dannenfelser RM. In vitro hemolysis: Guidance for the pharmaceutical scientist. J Pharm Sci. 2006;95(6):1173–1176. doi: 10.1002/jps.20627.

39 De Sena Pereira VS, Silva de Oliveira CB, Fumagalli F, da Silva Emery F, da Silva NB, de Andrade-Neto VF. Cytotoxicity, Hemolysis and In Vivo Acute Toxicity of 2-Hydroxy-3-Anilino-1,4-Naphthoquinone Derivatives. Toxicol Rep. 2016;3:756–762. doi: 10.1016/j.toxrep.2016.09.007.

40 Abdelaziz AA, Kamer AMA, Al-Monofy KB, et al. A purified and lyophilized Pseudomonas aeruginosa derived pyocyanin induces promising apoptotic and necrotic activities against MCF-7 human breast adenocarcinoma. Microb Cell Fact. 2022;21(1):262. doi: 10.1186/s12934-022-01988-x.

41 Gkinali AA, Matsakidou A, Paraskevopoulou A. Characterization of Tenebrio molitor Larvae Protein Preparations Obtained by Different Extraction Approaches. Foods. 2022;11(23):3852. doi: 10.3390/foods11233852.

42 Lidor O, Al-Quntar A, Pesci EC, Steinberg D. Mechanistic analysis of a synthetic inhibitor of the Pseudomonas aeruginosa LasI quorum-sensing signal synthase. Sci Rep. 2015;5:16569. doi: 10.1038/srep16569.

43 Song W, Tian Z, Wang Y, Yin Y, Zhang H, Xu C, et al. Isoliquiritigenin targets las, rhl, and Pqs quorum sensing systems to mitigate the virulence and infection of Pseudomonas aeruginosa. BMC Microbiol. 2025;25(1):[número de artículo o páginas si aplica]. doi: 10.1186/s12866-025-04298-5.

44 Ahmadzai A, Amirkhezi F, Bayan AM, Adel MQ, Sahar A, Mowahhed MR, et al. Mechanistic insights from docking and dynamics: soy isoflavonoids disrupt Pseudomonas aeruginosa quorum sensing by targeting AHL synthase LasI. Afghan J Basic Med Sci. 2026;3(1):53–74. doi: 10.62134/khatamuni.145.

45 Careviæ Miliæeviæ T, Novoviæ K, Stojkoviæ D, Kostiæ M, Kritsi E, Zoumpoulakis P, et al. Eriodictyol and diosmetin protective potential in skin infection: antimicrobial action, gene and molecular targets, and keratinocyte protection against bacteria-induced damage. Arch Pharm (Weinheim). 2025;358(7):e70047. doi: 10.1002/ardp.70047.

46 Bottomley MJ, Muraglia E, Bazzo R, Carfì A. Molecular insights into quorum sensing in the human pathogen Pseudomonas aeruginosa from the structure of the virulence regulator LasR bound to its autoinducer. J Biol Chem. 2007;282(18):13592–13600. doi: 10.1074/jbc.M700556200.

47 Amin A, Hanif M, Abbas K, Ramzan M, Rasheed A, Zaman A, et al. Studies on effects of umbelliferone derivatives against periodontal bacteria: antibiofilm, inhibition of quorum sensing and molecular docking analysis. Microb Pathog. 2020;144:104184. doi: 10.1016/j.micpath.2020.104184.

48 Bera AK, Atanasova V, Robinson H, Eisenstein E, Coleman JP, Pesci EC, et al. Structure of PqsD, a Pseudomonas quinolone signal biosynthetic enzyme, in complex with anthranilate. Biochemistry. 2009;48(36):8644–8654. doi: 10.1021/bi9009055.

49 Feng X, Shang J, Gu Z, Gong J, Chen Y, Liu Y. Azelaic acid: mechanisms of action and clinical applications. Clin Cosmet Investig Dermatol. 2024;17:2359–2371. doi: 10.2147/CCID.S485237.

50 Sauer N, Oœlizo M, Brzostek M, Wolska J, Lubaszka K, Karłowicz-Bodalska K. The multiple uses of azelaic acid in dermatology: mechanism of action, preparations, and potential therapeutic applications. Postepy Dermatol Alergol. 2023;40(6):716–724. doi: 10.5114/ada.2023.133955.

51 Blaskovich MAT, Elliott AG, Kavanagh AM, Ramu S, Mangan M, Huang JX, et al. In vitro antimicrobial activity of acne drugs against skin-associated bacteria. Sci Rep. 2019;9:14658. doi: 10.1038/s41598-019-50746-4.

52 Charnock C, Brudeli B, Klaveness J. Evaluation of the antibacterial efficacy of diesters of azelaic acid. Eur J Pharm Sci. 2004;21(4):589–596. doi: 10.1016/j.ejps.2003.12.006.

53 Arpa MD, Biltekin Kaleli S, Doğan N. Hydroxypropyl-β-cyclodextrin-enhanced azelaic acid hydrogel for acne treatment: evaluation of antimicrobial, anti-inflammatory, and skin penetration properties. J Pharm Innov. 2025;20:106. doi: 10.1007/s12247-025-10020-9.

54 Bandzerewicz A, Herman A, Dutkowska E, Niebuda K, Ruśkowski P, Gadomska-Gajadhur A. Combining polyesters of citric and azelaic acids to obtain potential topical application biomaterials with antimicrobial activity. Front Bioeng Biotechnol. 2025;13:1579630. doi: 10.3389/fbioe.2025.1579630.

55 Petrovici AG, Spennato M, Bîtcan I, Péter F, Cotarcă L, Todea A, et al. A comprehensive review of azelaic acid pharmacological properties, clinical applications, and innovative topical formulations. Pharmaceuticals (Basel). 2025;18(9):1273. doi: 10.3390/ph18091273.

56 Mariano-Rodriguez C, Nava-Martinez P, Diaz-Molina VL. Azelaic acid in dermatology: a review of its mechanism of action. Cureus. 2025;17(10):e94491. doi: 10.7759/cureus.94491.

57 Matijaševiæ D, et al. Inhibition of Pseudomonas aeruginosa quorum sensing-regulated behaviors by mushroom extracts. J Ethnopharmacol. 2025;353:120466. doi: 10.1016/j.jep.2025.120466.

58 Vadakkan K, Ngangbam AK, Sathishkumar K, Rumjit NP, Cheruvathur MK. A review of chemical signaling pathways in the quorum sensing circuit of Pseudomonas aeruginosa. Int J Biol Macromol. 2024. doi: 10.1016/j.ijbiomac.2023.127861.

59 Li WR, Zeng TH, Xie XB, Shi QS, Li CL. Inhibition of the pqsABCDE and pqsH in the pqs quorum sensing system and related virulence factors of Pseudomonas aeruginosa PAO1 strain by farnesol. Int Biodeterior Biodegradation. 2020;151:104956. doi: 10.1016/j.ibiod.2020.104956.

60 Liu J, Wang Z, Zeng Y, Wang W, Tang S, Jia A. 1H-pyrrole-2,5-dicarboxylic acid, a quorum sensing inhibitor from an endophytic fungus in Areca catechu L., acts as antibiotic accelerant against Pseudomonas aeruginosa. Front Cell Infect Microbiol. 2024;14:1413728. doi: 10.3389/fcimb.2024.1413728.

61 Jabłoñska-Wawrzycka A, Rogala P, Czerwonka G, Michałkiewicz S, Hodorowicz M, Gałczyñska K, et al. Tuning anti-biofilm activity of manganese(II) complexes: linking biological effectiveness of heteroaromatic complexes of alcohol, aldehyde, ketone, and carboxylic acid with structural effects and redox activity. Int J Mol Sci. 2021;22(9):4847. doi: 10.3390/ijms22094847.

62 Diaz-Guerrero M, López-Jácome LE, Franco-Cendejas R, Coria-Jiménez R, Martínez-Zavaleta MG, González-Pedrajo B, et al. Curcumin inhibits type III secretion of Pseudomonas aeruginosa. PeerJ. 2025;13:e19725. doi: 10.7717/peerj.19725.

63 González GM, Contreras RG, Alvarado-Salazar JA, García Valdés I, Valdés NG, Lugo Gutiérrez DA, et al. Effect of curcumin and 5-bromoindole-3-carboxaldehyde on the inhibition of quorum sensing-dependent virulence factors in Pseudomonas aeruginosa. bioRxiv [Preprint]. 2024. doi: 10.1101/2024.07.29.605599.

64 García-Contreras R, Martínez-Vázquez M, Velázquez Guadarrama N, Villegas Pañeda AG, Hashimoto T, Maeda T, et al. Resistance to the quorum-quenching compounds brominated furanone C-30 and 5-fluorouracil in Pseudomonas aeruginosa clinical isolates. Pathog Dis. 2013;68(1):8–11. doi: 10.1111/2049-632X.12039.

65 Abbas HA, Shaldam MA, Eldamasi D. Inhibition of quorum sensing in Pseudomonas aeruginosa by sitagliptin. Curr Microbiol. 2020;77(6):1051–1060. doi: 10.1007/s00284-020-01909-4.

66 Kalia M, Singh PK, Yadav VK, Yadav BS, Sharma D, Narvi SS, et al. Structure-based virtual screening for identification of potential quorum sensing inhibitors against LasR master regulator in Pseudomonas aeruginosa. Microb Pathog. 2017;107:136–143. doi: 10.1016/j.micpath.2017.03.026.

67 Anju VT, Busi S, Mohan MS, Ranganathan S, Ampasala DR, Kumavath R, et al. In vivo, in vitro and molecular docking studies reveal the anti-virulence property of hispidulin against Pseudomonas aeruginosa through modulation of quorum sensing. Int Biodeterior Biodegradation. 2022;174:105487. doi: 10.1016/j.ibiod.2022.105487.

68 Siam MH, Sirajee AS, Limon MH, Hossain MA, Sultana M. In silico identification of quorum sensing inhibitors against LasR protein in a clinical isolate of multidrug-resistant Pseudomonas aeruginosa DMC-27b. F1000Res. 2024;13:62. doi: 10.12688/f1000research.131728.1.

69 Zhou Z, Ma S. Recent advances in the discovery of PqsD inhibitors as antimicrobial agents. ChemMedChem. 2017;12(6):420–425. doi: 10.1002/cmdc.201700015.

70 El-Mowafy SA, Abd El Galil KH, Habib EE, Shaaban MI. Quorum sensing inhibitory activity of sub-inhibitory concentrations of β-lactams. Afr Health Sci. 2017;17(1):199–207. doi: 10.4314/ahs.v17i1.25.

71 Kalia VC, Wood TK, Kumar P. Evolution of resistance to quorum-sensing inhibitors. Microb Ecol. 2014;68(1):13–23. doi: 10.1007/s00248-013-0316-y.

72 Siam MH, Sirajee AS, Limon MH, et al. In silico identification of quorum sensing inhibitors against LasR protein in a clinical isolate of multidrug resistant Pseudomonas aeruginosa DMC-27b. F1000Res. 2024;13:62. doi: 10.12688/f1000research.131728.1.

73 Goel N, Ghosh M, Jain D, Sinha R, Khare SK. Inhibition and eradication of Pseudomonas aeruginosa biofilms by secondary metabolites of Nocardiopsis lucentensis EMB25. RSC Med Chem. 2023;14(4):745–756. doi: 10.1039/d2md00439a.

74 Dergez Á, et al. Inhibition of exopolysaccharide biopolymer and pyocyanin virulence factors produced by Pseudomonas aeruginosa 1604 by salicylic compounds. Period Polytech Chem Eng. 2014;58(1):75–80. doi: 10.3311/PPch.7207.

75 Noto MJ, Burns WJ, Beavers WN, Skaar EP. Mechanisms of Pyocyanin Toxicity and Genetic Determinants of Resistance in Staphylococcus aureus. J Bacteriol. 2017;199(17):e00221–17. doi: 10.1128/JB.00221-17.

76 Barton TE, et al. Galleria mellonella as an Antimicrobial Screening Model. JoVE. 2024;(212):e67210. doi: 10.3791/67210.

77 Ramakrishnan R, Daly AC. Revisiting ISO 10993-5 In Vitro Cytotoxicity Standard Tests for Evaluation of Extracellular Matrix-Based Biomaterials. Int J Biomater. 2026;2026:7395612. doi: 10.1155/ijbm/7395612.

78 Elizondo-Luevano JH, Quintanilla-Licea R, Castillo-Hernández SL, Sánchez-García E, Bautista-Villarreal M, González-Meza GM, et al. In Vitro Evaluation of Anti-Hemolytic and Cytotoxic Effects of Traditional Mexican Medicinal Plant Extracts on Human Erythrocytes and Cell Cultures. Life (Basel). 2024;14(9):1176. doi: 10.3390/life14091176.

79 Sæbø IP, Bjørås M, Franzyk H, Helgesen E, Booth JA. Optimization of the Hemolysis Assay for the Assessment of Cytotoxicity. Int J Mol Sci. 2023;24(3):2914. doi: 10.3390/ijms24032914.

80 Jeong H, Hwang J, Lee H, et al. In vitro blood cell viability profiling of polymers used in molecular assembly. Sci Rep. 2017;7(1):9481. doi: 10.1038/s41598-017-10169-5.

